# Glycolytic flux-signaling controls mouse embryo mesoderm development

**DOI:** 10.1101/2021.12.20.473441

**Authors:** Hidenobu Miyazawa, Marteinn T. Snaebjornsson, Nicole Prior, Eleni Kafkia, Henrik M. Hammarén, Nobuko Tsuchida-Straeten, Kiran R. Patil, Martin Beck, Alexander Aulehla

## Abstract

How cellular metabolic state impacts cellular programs is a fundamental, unresolved question. Here we investigated how glycolytic flux impacts embryonic development, using presomitic mesoderm (PSM) patterning as the experimental model. First, we identified fructose 1,6-bisphosphate (FBP) as an *in vivo* sentinel metabolite that mirrors glycolytic flux within PSM cells of post-implantation mouse embryos. We found that medium-supplementation with FBP, but not with other glycolytic metabolites, such as fructose 6-phosphate and 3-phosphoglycerate, impaired mesoderm segmentation. To genetically manipulate glycolytic flux and FBP levels, we generated a mouse model enabling the conditional overexpression of dominant active, cytoplasmic Pfkfb3 (cytoPfkfb3). Overexpression of cytoPfkfb3 indeed led to increased glycolytic flux/FBP levels and caused an impairment of mesoderm segmentation, paralleled by the downregulation of Wnt-signaling, reminiscent of the effects seen upon FBP-supplementation. To probe for mechanisms underlying glycolytic flux-signaling, we performed subcellular proteome analysis and revealed that cytoPfkfb3 overexpression altered subcellular localization of certain proteins, including glycolytic enzymes, in PSM cells. Specifically, we revealed that FBP supplementation caused depletion of Pfkl and Aldoa from the nuclear-soluble fraction. Combined, we propose that FBP functions as a flux-signaling metabolite connecting glycolysis and PSM patterning, potentially through modulating subcellular protein localization.

## Introduction

Living systems have the critical ability to sense environmental cues, and to integrate this information with cellular functions by modulating their metabolic activity (1, 2). The changes in metabolic activity, in turn, are sensed by multiple mechanisms to ensure that metabolic state matches cellular demands. Such mechanisms, referred to as metabolite sensing and signaling, generally consist of ‘sentinel metabolites’ and ‘sensor molecules’ (3, 4). Sentinel metabolites mirror nutrient availability or cellular metabolic state by their levels. These metabolites can, in addition, potentially induce cellular responses, if their levels are linked to the activity of sensor molecules, such as proteins and RNAs. Well known examples of metabolite sensing and signaling include the mechanistic target of rapamycin (mTOR), which responds to altered levels of amino acids and couples nutritional availability with cell growth (5), or AMP-activated protein kinase (AMPK), which senses adenosine monophosphate (AMP) levels and ensures that cellular bioenergetic demand matches cellular energetic state (6).

Importantly, the role of metabolite signaling is not limited to detecting nutrient availability to match metabolic activity and cellular demands. Recent work has highlighted the emerging link between central carbon metabolism and other cellular programs, such as gene regulation. For instance, by controlling the abundance of rate-limiting substrates used for post-translational modificiations, such as acetyl-CoA, metabolic activity can directly impact gene expression (7–9). Glycolytic metabolites can also serve as signaling molecules that impact signal transduction directly. In yeast, for example, the glycolytic metabolite fructose 1,6-bisphosphate (FBP) has been shown to regulate the pro-proliferative RAS signaling cascade by interacting with the guanine nucleotide exchange factor Sos1 (10). Notably, the connection between metabolic activity and other cellular programs can also occur at the level of metabolic enzymes with non-canonical, moonlighting functions (9, 11, 12). In situations when moonlighting and canonical enzyme function are inter-dependent, a direct link between cellular metabolic state and moonlighting function is established. One such example is the glycolytic enzyme glyceraldehyde 3-phosphate dehydrogenase (Gapdh), which moonlights as an RNA-binding protein regulating translation when not engaged in its glycolytic function (13). While these studies highlight an intricate link between central carbon metabolism and other cellular functions, knowledge of metabolite signaling in more complex physiological settings, such as embryonic development, is still limited.

There are both classic (14) as well as more recent findings (9, 15–21) indicating that glucose metabolism and developmental programs are indeed linked. For instance, in mouse and chick embryos, the presomitic mesoderm (PSM) shows intrinsic differences in the expression levels of glycolytic enzymes, leading to the establishment of a glycolytic activity gradient along the anterior-posterior axis (15, 16). The key question that remains largely unanswered is how a change in cellular metabolic activity is sensed and mechanistically linked to developmental programs. To address this fundamental question, we focused on mouse embryos at the organogenesis stage following gastrulation, when glucose metabolism is rewired dynamically in time and space in response to extrinsic environmental cues and intrinsic developmental programs (15–17). At this stage, the PSM is periodically segmented into somites, the precursors of vertebrae and skeletal muscles in vertebrates (22). PSM patterning and somite formation is controlled by the Wnt, FGF, and retinoic acid-signaling pathways, which show a graded activity along the anterior-posterior axis. In addition, PSM segmentation is linked to a molecular oscillator, the segmentation clock, comprised of several, interconnected signaling pathways (Notch, Wnt, Fgf) that show rhythmic activation cycles in PSM cells, with a period matching the rate of somite formation, e.g. ∼2 hours in mouse embryos (23–29). The interplay between graded and oscillatory signaling dynamics within the PSM controls somite formation in time and space. Previously, a link between glycolytic activity and graded signaling activities has been found (15, 16, 30). In particular, evidence was found that glycolysis is part of a feedback loop linking (graded) FGF- and Wnt-signaling pathway activities (16, 30). Although these studies revealed a link between glycolysis and morphogen signaling during PSM patterning, it remains unclear how a change in glycolytic activity is sensed and mechanistically linked to signaling. In this study we aimed to identify the relevant sentinel glycolytic metabolite(s), its function, and its sensing mechanism during mouse mesoderm development, using genetics, metabolomics, and proteomics approaches.

## Results

### Steady state levels of FBP mirror glycolytic flux within PSM cells

In order to identify sentinel metabolites whose levels reflect glycolytic-flux within PSM cells, we quantified steady state metabolite levels in PSM samples cultured in a range of glucose concentrations (0.5 mM to 10 mM glucose). We first verified that higher glucose concentrations led to higher glycolytic activity in PSM cells using lactate secretion as a proxy (Figure 1A). Also, we analyzed somite formation and PSM patterning at different glucose concentrations (Figure S1). Using real-time imaging of the segmentation clock as a dynamic readout, we found ongoing periodic morphological segmentation, axis elongation, and oscillatory clock activity throughout the PSM. Segmentation proceeded normally, at least qualitatively, at these glucose concentrations. We then used this experimental setting to analyze steady state levels of metabolites in central carbon metabolism by gas chromatography mass spectrometry (GC-MS), following three-hour incubation of PSM explants at glucose concentrations ranging from 0.5 mM to 10 mM. Amongst the 57 metabolites quantified, 14 metabolites showed significant linear correlation (*p*-value < 0.01) with extracellular glucose levels (Figure 1B). In particular, fructose 1,6-bisphosphate (FBP) showed the highest positive linear relationship (Pearson correlation coefficient = 0.99) with extracellular glucose availability. Notably, unlike secreted lactate levels, the linear relationship was maintained across all glucose concentrations examined (Figure 1A). In addition, FBP showed the highest fold-change response to glucose titration with a 45-fold increase in its levels from 0.5 mM to 10 mM glucose (Figure 1A, 1C). These results show that steady state FBP levels tightly reflect glucose availability, indicating the potential of FBP to serve as a sentinel metabolite for glycolytic-flux within PSM cells.

**Fig. 1.**
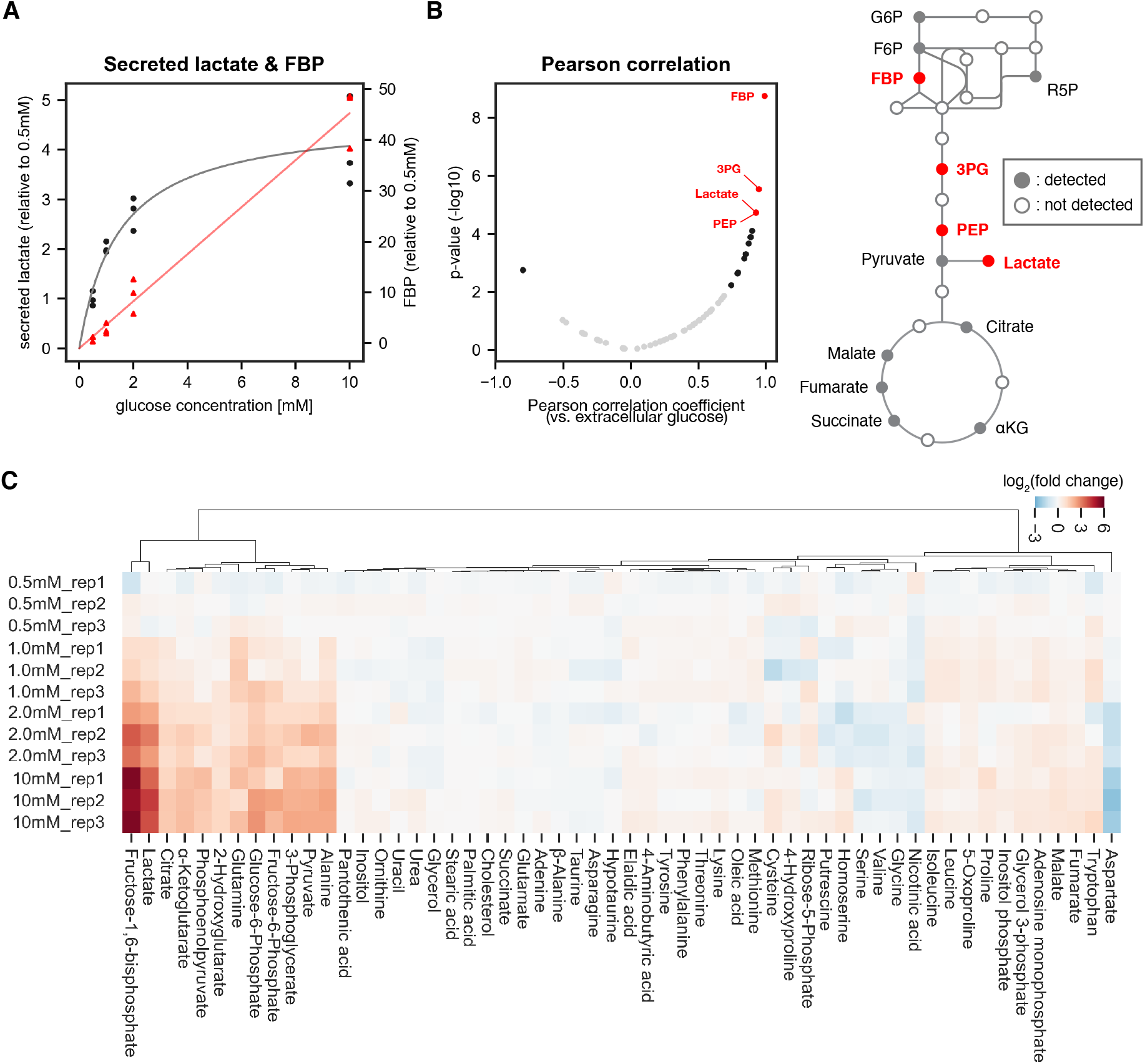
Identifying sentinel metabolites that mirror glycolytic flux. The amount of secreted lactate and intracellular metabolites within PSM explants were measured by gas chromatography mass spectrometry (GC-MS; n = 3 biological replicates for each condition). The explants were cultured for three hours *ex vivo* in 0.5 mM, 1.0 mM, 2.0 mM, or 10 mM glucose. **(A)** The relative amount of secreted lactate (shown in black circles) and intracellular fructose 1,6-bisphosphate (FBP; shown in red triangles) under various glucose conditions. The gray and red lines show the Michaelis-Menten fit (Vmax = 4.7 arbitrary unit, Km = 1.5 mM) for secreted lactate and the linear regression line for intracellular FBP, respectively. **(B)** Pearson correlation analysis between intracellular metabolite levels and extracellular glucose levels. Metabolites showing significant correlation (*p*-value < 0.01) are shown in black. Those with a |Pearson correlation coefficient| > 0.9 are highlighted in red. Abbreviations: G6P, glucose 6-phosphate; F6P, fructose 6-phosphate; R5P, ribose 5-phosphate; FBP, fructose 1,6-bisphosphate; 3PG, 3-phosphoglycerate; PEP, phosphoenol pyruvate; αKG, α-ketoglutarate. **(C)** Hierarchical clustering heatmap of metabolites detected in the PSM explants. Fold changes were calculated using 0.5 mM glucose condition as the reference. Hierarchical clustering was performed using Ward’s method with Euclidean distance.

### Altered mesoderm development caused specifically by FBP supplementation

To test for a potential functional role of those sentinel metabolites that we identified, we next performed medium-supplementing experiments with the goal of altering intracellular metabolite levels. To this end, we supplemented the control culture medium with high levels of either fructose 6-phosphate (F6P), FBP, or 3-phosphoglycerate (3PG) and scored the effect at the level of morphological segment formation, elongation, and also oscillatory segmentation clock activity, using real-time imaging quantifications. Interestingly, FBP supplementation impaired mesoderm segmentation and elongation and disrupted segmentation clock activity in the posterior PSM (Figure 2A, S2A, S2B). In contrast, glycolytic metabolites upstream (*i*.*e*. F6P) or downstream (*i*.*e*. 3PG, pyruvate) of FBP did not cause such effects (Figure 2A, S2A, S2B ; the effect of pyruvate supplementation was described in (15)). We also tested the effect of FBP supplementation on gene expression, focusing on an FGF-target gene *Dusp4* (31) and a Wnt-target gene *Msgn* (32). Supplementation of FBP, but not F6P, caused a downregulation of *Dusp4* and *Msgn* mRNA expression in a dose-dependent manner (Figure 2B), accompanying reduction of mesoderm segmentation and elongation (Figure 2C, 2D). Of note, at intermediate concentration (10 mM) of FBP supplementation, only the Wnt-taget gene *Msgn* was downregulated, while the Fgf-target gene *Dusp4* showed expression comparable to control samples, indicating potential dose-specific effects of FBP. To validate the effects seen upon exogenous addition of FBP, we investigated the uptake of FBP by stable isotope (^13^C) tracing. We cultured PSM explants in medium supplemented with fully ^13^C-labelled FBP (^13^C_6_-FBP) and analyzed ^13^C-labelling of intracellular metabolites by liquid chromatography mass spectrometry (LC-MS). Following three hours of incubation with ^13^C_6_-FBP, ^13^C-labeling was detected in glycolytic intermediates downstream of FBP (Figure S2C), confirming the uptake of labeled carbons by the explants. Since we also detected that a small fraction of ^13^C_6_-FBP broke down to ^13^C_6_-fructose monophosphate (F6P and/or fructose 1-phosphate (F1P)) in the culture medium during incubation (data not shown), we performed additional control experiments by culturing PSM explants in F1P-supplemented medium. Similar to F6P, supplementation of F1P did not cause any detectable phenotype at the level of segmentation clock activity or elongation (Figure S2D).

**Fig. 2.**
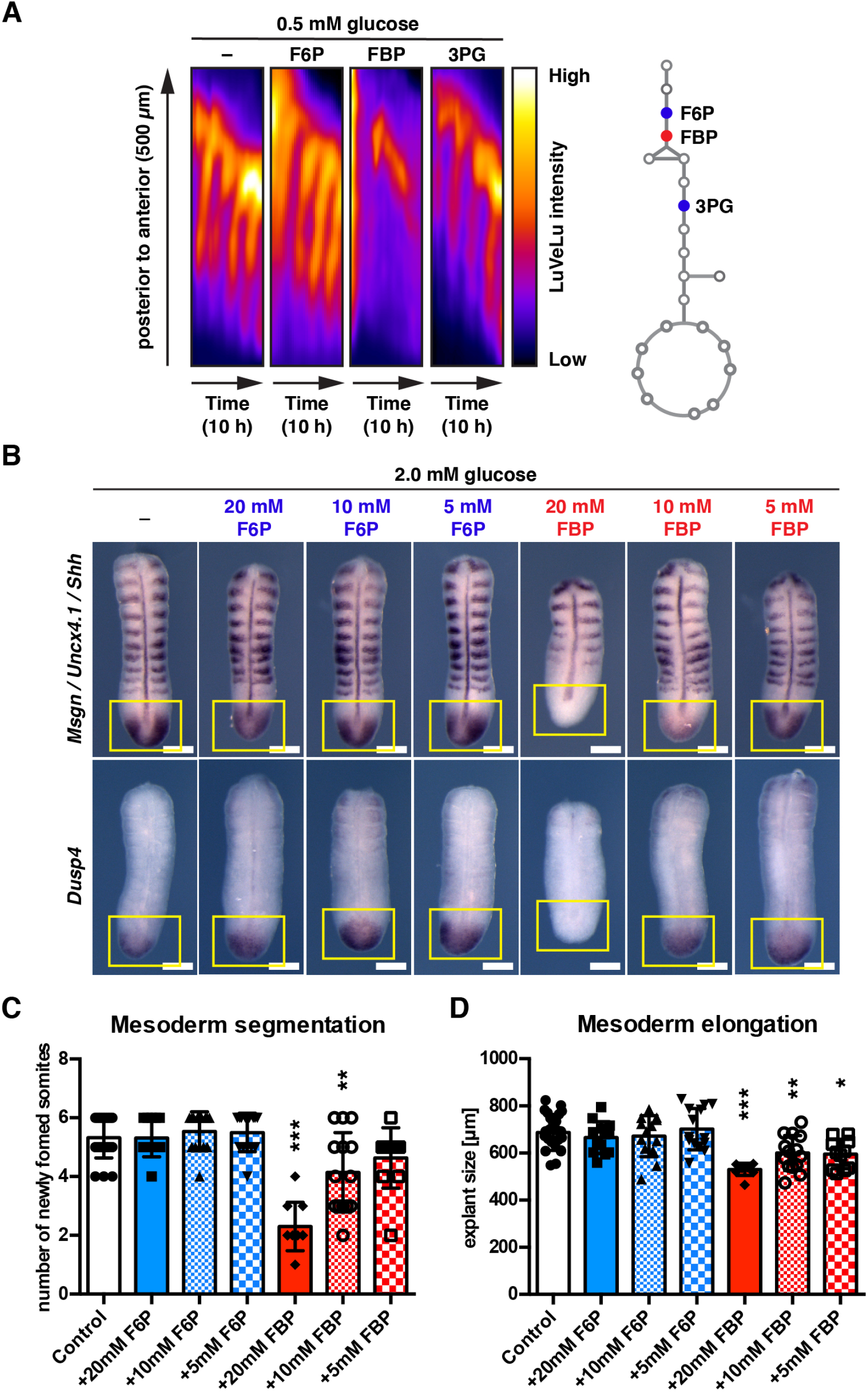
FBP supplementation impacts mesoderm segmentation and elongation in a dose-dependent manner. **(A)** Kymographs showing dynamics of the Notch signaling activity reporter LuVeLu in PSM explants treated with 20 mM of the indicated metabolite. **(B)** Whole mount *in situ* hybridization analysis for the FGF (*i*.*e. Dusp4*) and the Wnt target (*i*.*e. Msgn*) gene expression in the PSM. PSM explants were incubated for 12 hours in the presence of F6P or FBP. Expression domains of *Dusp4* and *Msgn* are indicated by yellow squares. *Shh* and *Uncx4*.*1* were used as a marker for the neural tissue and posterior somite boundary, respectively. Scale bar, 200 µm. **(C, D)** The number of newly formed somites (C) and the length of PSM explants (D) after 12-hour *ex vivo* culture (one-way ANOVA with Tukey’s post-hoc test, \**p*-value < 0.05, \*\**p*-value < 0.01, \*\*\**p*-value < 0.001 versus control).

As a related finding, we observed that upon glucose titration, the expression of Wnt-signaling target genes in PSM explants is anti-correlated with glucose availabilty/glycolytic activity: while lowering glucose concentration (from 5.0 mM to 0.5 mM) correlated with an upregulation of several Wnt target genes, such as *Axin2, Ccnd1*, and *Myc*, the opposite effect was found when glucose concentration was increased (from 5.0 mM to 25 mM) (Figure S3).

Combined, our findings hence suggest that FBP, but not other glycolytic intermediates such as F6P, F1P or 3PG, is a flux-sentinel and signaling metabolite, as it impacts mesoderm development and gene expression in a dose-dependent manner.

### Generating a conditional *cytoPfkfb3* transgenic mouse line as a genetic tool to increase glycolysis

Our findings thus far show that intracellular FBP levels respond dynamically to an alteration in glycolytic flux (Figure 1), and importantly, that FBP, but not its precursor metabolite F6P, impacts PSM development in a dose-dependent manner (Figure 2). Based on these observations, we next sought a way to manipulate glycolytic flux at the level of the phosphofructokinase (Pfk) reaction and importantly, in a genetic manner (Figure 3A). Pfk converts F6P into FBP, the first committed step in glycolysis, and plays a critical role in regulating glycolytic flux (33, 34). We generated transgenic mice enabling conditional overexpression of a mutant Pfkfb3 [*i*.*e*. Pfkfb3(K472A/K473A) (35)]. Pfkfb3 generates fructose 2,6-bisphosphate (F2,6BP), a potent allosteric activator of Pfk (Figure 3A). A previous study showed that Pfkfb3(K472A/K473A) localises exclusively to the cytoplasm, and that this cytoplasmically-localized Pfkfb3 (hereafter termed as cytoPfkfb3) activates glycolysis (35). Indeed, in PSM explants from transgenic embryos with ubiquitous overexpression of cytoPfkfb3, we found increased glycolysis based on the analysis of lactate secretion (Figure 3B). In addition, we found that in *cytoPfkfb3* embryos, lactate secretion changed in a glucose-dose dependent manner (Figure 3B). Next we investigated steady state metabolite levels in control and transgenic PSM explants cultured in 10 mM glucose condition. Among the 57 metabolites quantified by GC-MS, FBP and lactate were significantly increased in transgenic PSM explants, while aspartate, glucose 6-phosphate, and glutamate were significantly decreased (Figure 3C, S4). These findings mirrored the results in wild-type PSM explants upon glucose titration (Figure 1). We hence conclude, that the overexpression of cytoPfkfb3 leads to activation of glycolysis at the level of Pfk. More generally, the *cytoPfkfb3* transgenic mouse line represents a potentially powerful new genetic model to study the role of glycolytic activation.

**Fig. 3.**
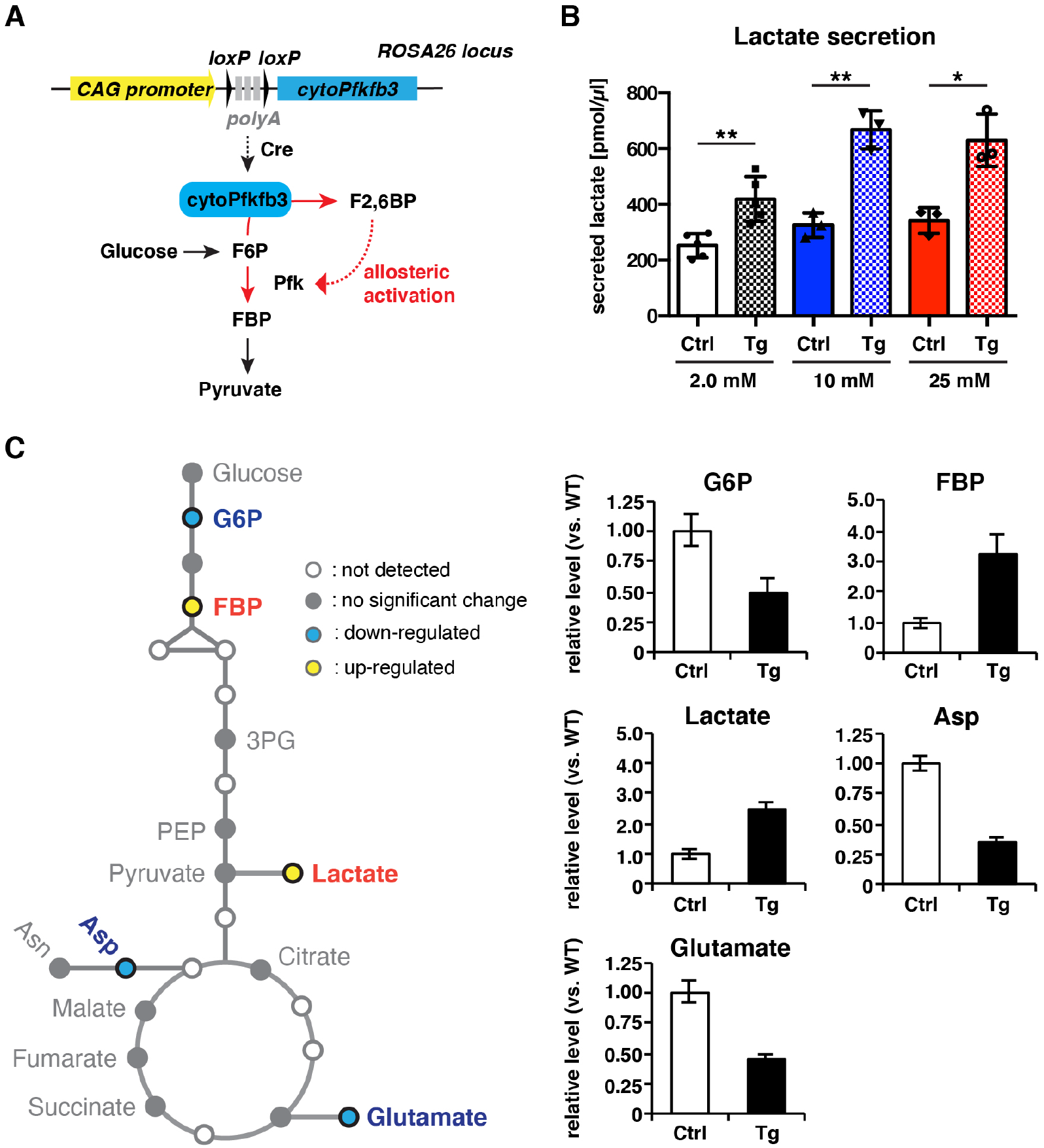
cytoPfkfb3 overexpression causes an increase in glycolytic flux and FBP levels within PSM cells. **(A)** Conditional *cytoPfkfb3* transgenic mice were generated to activate glycolysis through allosteric activation of Pfk. **(B)** Quantification of secreted lactate in control and *cytoPfkfb3* transgenic PSM explants cultured 12 hours under varying concentrations of glucose (unpaired Welch’s *t* -test, \**p*-value < 0.05, \*\**p*-value < 0.01). **(C)** Measurement of steady state metabolite levels by GC-MS (n = 4 biological replicates for each condition) in control (Ctrl) and *cytoPFKfb3* (Tg; crossed to *Hprt-Cre* line) explants cultured for three hours in medium containing 10 mM glucose. SAM (Significance Analysis for Microarrays) analysis was performed using a significance threshold *δ* = 0.9, which corresponds to a false discovery rate (FDR) = 0.012. G6P, glucose 6-phosphate. FBP, fructose 1,6-bisphosphate. 3PG, 3-phosphoglycerate. PEP, phosphoenol pyruvate. Asp, aspartate. Asn, asparagin.

### Functional consequence of cytoPfkfb3 overexpression on PSM development

We then investigated the functional consequences of cytoPfkfb3 overexpression on mesoderm development. Constitutive overexpression of cytoPfkfb3 from fertilization caused embryonic lethality, as no transgenic pups were recovered (n = 30 pups, N = 6 litters). We have not yet investigated the precise timepoint and cause of lethality. At embryonic day 10.5 (E10.5), cytoPfkfb3 transgenic embryos were morphologically indistinguishable from their littermates, but had slightly fewer somites (Figure 4A; control: 38 ± 1.5 somites, transgenic: 35 ± 3.9 somites).

**Fig. 4.**
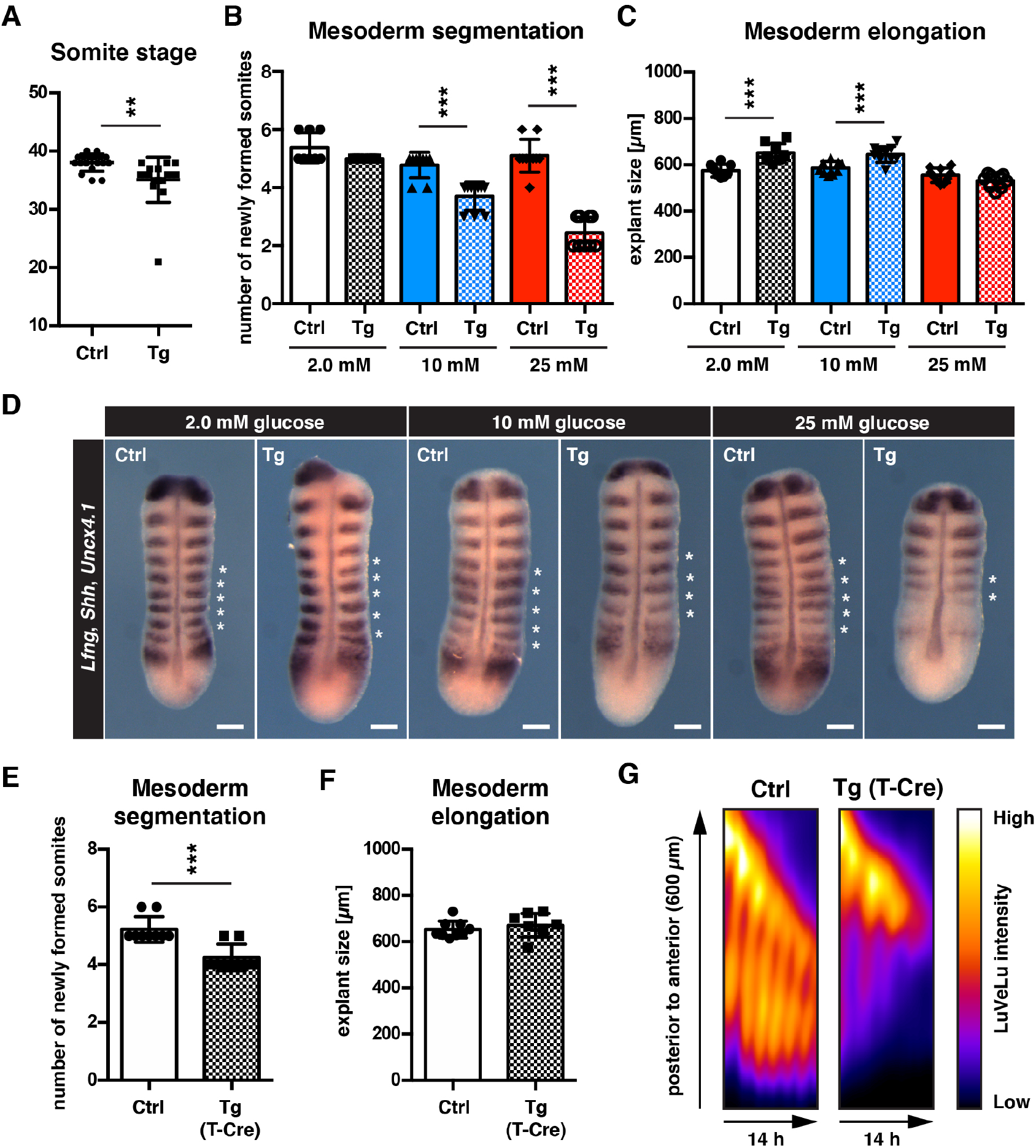
cytoPfkfb3 overexpression impacts mesoderm development in a glucose-concentration dependent manner. **(A)** Total number of somites in E10.5 embryos (mean ± s.d; unpaired Welch’s *t* -test; \*\**p*-value < 0.01). Ctrl, control embryos; Tg, *cytoPfkfb3* embryos (crossed to *Hprt-Cre* line). **(B**,**C)** Number of formed somites and quantification of PSM explant length after 12-hour *in vitro* culture. **(D)** Whole mount mRNA *in situ* hybridization analysis for *Lfng, Shh*, and *Uncx4*.*1* in PSM explants after 12-hour *in vitro* culture at varying glucose concentrations. Asterisks denote somites that formed during the *in vitro* culture. Scale bar, 100 µm. **(E**,**F)** Effect of mesoderm-specific overexpression of cytoPfkfb3 on PSM segmentation and elongation (12-hour incubation). The PSM explants were cultured in medium containing 10 mM glucose. Bar graphs show the number of newly formed somites during the culture (E), and the length of explants after the culture (F; mean ± s.d; unpaired Welch’s *t* -test; \*\*\**p*-value < 0.001). **(G)** Real-time quantification of segmentation clock activity using Notch signaling activity reporter LuVeLu in PSM explants, shown as kymographs. Note that oscillatory reporter activity ceased in cytoPfkfb3/T-Cre samples during the experiment, while control samples showed ongoing periodic activity.

To analyze the impact of cytoPfkfb3 overexpression on mesoderm development in a more dynamic and quantitative manner, we analyzed mesoderm segmentation, elongation and oscillatory clock activity in *cytoPfkfb3* and control explants cultured at various glucose concentrations. Consistent with our previous findings (Figure S1), control explants proceeded segmentation and PSM patterning in a qualitatively comparable manner, even when cultured at higher glucose concentrations (Figure 4B). In contrast, we found that somite formation was impaired in explants from *cytoPfkfb3* embryos in a glucose-dose dependent manner (Figure 4B). Overall growth during this 12-hour incubation seemed comparable or even increased in *cytoPfkfb3* transgenic explants, based on the size of explants after culture (Figure 4C). We also tested whether a mesoderm-specific cytoPfkfb3 overexpression has a similar effect on somite formation. Indeed, mesoderm specific cytoPfkfb3 overexpression, using Cre-expression driven by the promoter of the pan-mesoderm marker *Brachyury* (i.e. *T*-promoter-driven Cre (36)), showed similar reduction in segment formation, compared to control explants (Figure 4E, 4F). The real-time imaging quantification of segmentation clock activity revealed that in *cytoPfkfb3* explants cultured at 10 mM glucose, clock oscillations ceased after few cycles, in contrast to control samples (Figure 4G).

Molecularly, we found that the expression of the Wnt signaling target gene *Msgn* was downregulated in *cytoPfkfb3* explants, again in a glucose-concentration dependent manner. In contrast, we did not find an obvious change in the expression of *Dusp4*, an Fgf signaling target, which was maintained even at 25 mM glucose (Figure 5A, 5B). Taken together, these results show that cytoPfkfb3 overexpression, which we show leads to increased glycolytic flux, results in reduced segment formation, arrest of the segmentation clock oscillations and downregulation of Wnt-signaling target gene expression. In this regard, cytoPfkfb3 overexpression mirrors the results obtained with exogenous FBP-supplementation.

**Fig. 5.**
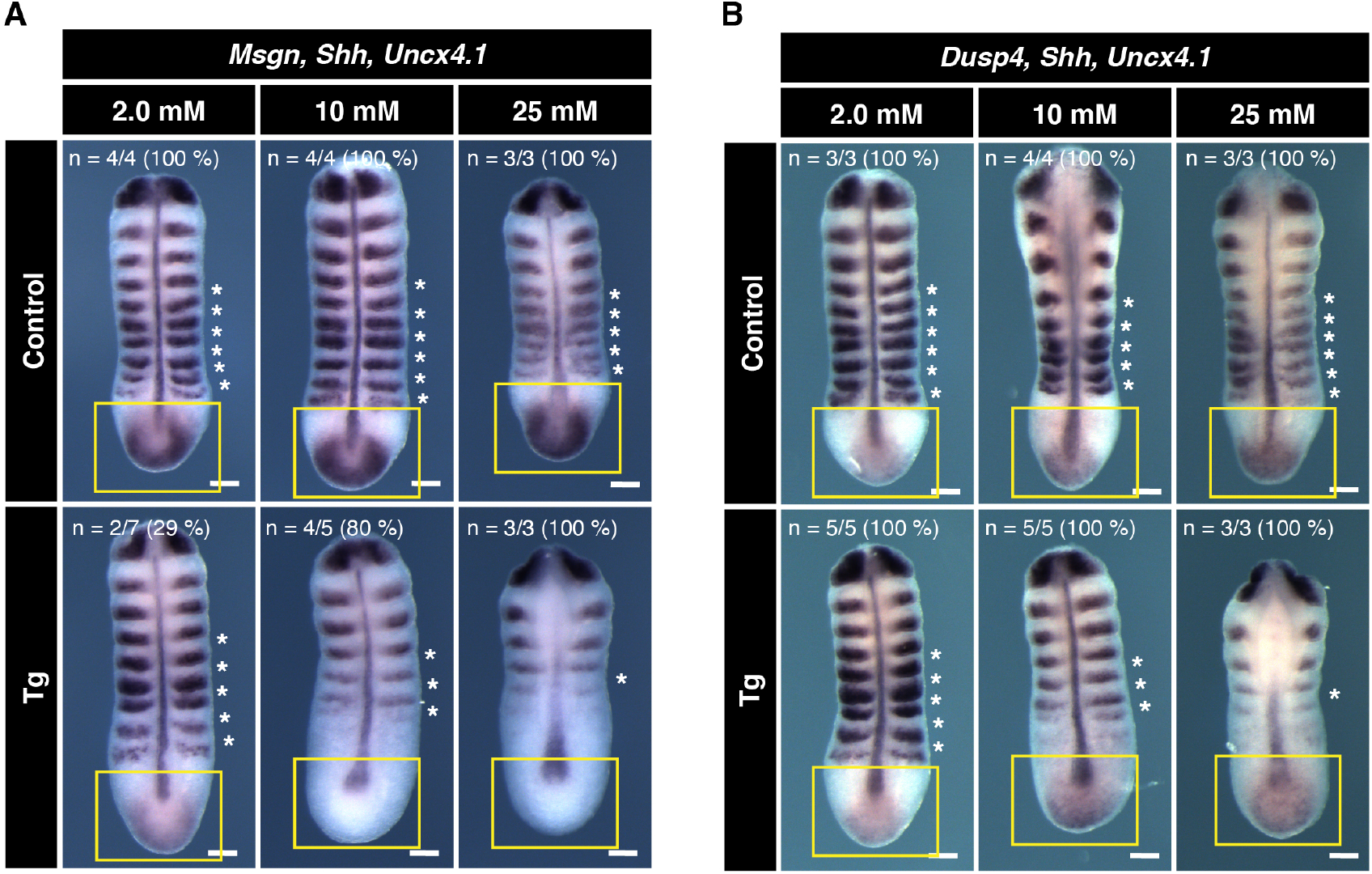
Effect of cytoPfkfb3 overexpression on Wnt and FGF target gene expression. **(A**,**B)** Whole mount mRNA *in situ* hybridization for *Msgn* (Wnt-target gene) and *Dusp4* (FGF-target gene) in the PSM explants. Explants were cultured for 12 hours under various glucose conditions, as indicated. *Shh* and *Uncx4*.*1* were used as a marker for neural tissue and posterior somite boundary, respectively. Expression domain of *Msgn* and *Dusp4* is indicated by yellow rectangles. Note the glucose-dose dependent loss of *Msgn* expression in *cytoPfkfb3* explants (Tg; crossed to *Hprt-Cre* line). In contrast, *Dusp4* expression appeared unaffected in cytoPfkfb3 explants. Asterisks mark somites that formed during the culture. Scale bar, 100 µm.

### cytoPfkfb3 overexpression impairs neural tube closure *ex vivo*

We also examined the effect of cytoPfkfb3 overexpression on other developmental events, *i*.*e*. neural tube closure (NTC). NTC is known to be vulnerable to glucose metabolism perturbation associated with maternal diabetes (37). In order to address the impact of cytoPfkfb3 overexpression on NTC, E8.5 cytoPfkfb3 embryos were cultured for 24 hours at normoglycemic conditions (50% rat serum / DMEM with 1.0 g/L glucose), using the whole embryo roller-culture (WEC) system (38). While all control embryos completed cranial NTC (n = 12/12) after 24-hours WEC, about 40% of the transgenic embryos (n = 7/18) failed to complete this process (Figure S5A–S5C). We also noticed a tendency that *cytoPfkfb3* embryos with neural tube defects (NTDs) had fewer somites (18 ± 1.6 somites) compared to wild-type (22 ± 2.2 somites) or the transgenic embryos that completed NTC (20 ± 1.6 somites) (Figure S5B). These results indicate that cytoPfkfb3 overexpression causes NTDs and a developmental delay when assessed in WEC. Interestingly, *cytoPfkfb3* transgenic embryos did not exhibit NTDs *in vivo*. We speculate this reflects a dependency of the NTD phenotype on environmental glucose concentrations, which are lower *in vivo* compared to WEC (39).

### Perturbation of glycolytic-flux and FBP levels alters subcellular localization of glycolytic enzymes

Our data thus far suggest that altered glycolysis, caused by either nutritional or genetic means, impairs PSM development, possibly mediated via the sentinel metabolite FBP. To probe for potential underlying mechanisms, we turned to the role of glycolytic enzymes. Interestingly, we had found that several glycolytic enzymes are localized in the nucleus in PSM cells, based on cell-fractionation analysis (Figure S6C, S6D). It had been proposed previously that the subcellular localization of glycolytic enzymes can change dynamically in response to altered glycolytic flux (40–42). We therefore aimed to systematically investigate the changes in subcellular protein localization in response to altered metabolic state in mouse embryos. To this end, we performed a proteome-wide cell-fractionation analysis in PSM explants cultured in various metabolic conditions.

Proteins were extracted from cytoplasmic, membrane, nuclear-soluble, chromatin-bound, and the remaining insoluble (labeled as ‘cytoskeletal’) fractions. We found that in samples cultured three hours in FBP-supplemented medium (and to a lesser extend in F6P-supplemented medium), proteins part of the glycolytic pathway (12 combined glycolytic enzymes) were reduced in the cytoskeletal and, to a lesser extent, the nuclear soluble fraction, relative to samples cultured in control medium. (Figure 6A, S6A, S6B). For several glycolytic enzymes detected in the nuclear soluble fraction, *i*.*e*. aldolase A (Aldoa), phosphofructokinase L (Pfkl), glyceraldehyde 3-phosphate dehydrogenase (Gapdh) and pyruvate kinase M (Pkm) (Figure S6E), we performed a targeted analysis using Western blotting. Interestingly, we found that amongst those tested enzymes, Aldoa and Pfkl were significantly depleted from the nuclear soluble fraction upon incubation in FBP-supplemented medium (Figure 6B).

**Fig. 6.**
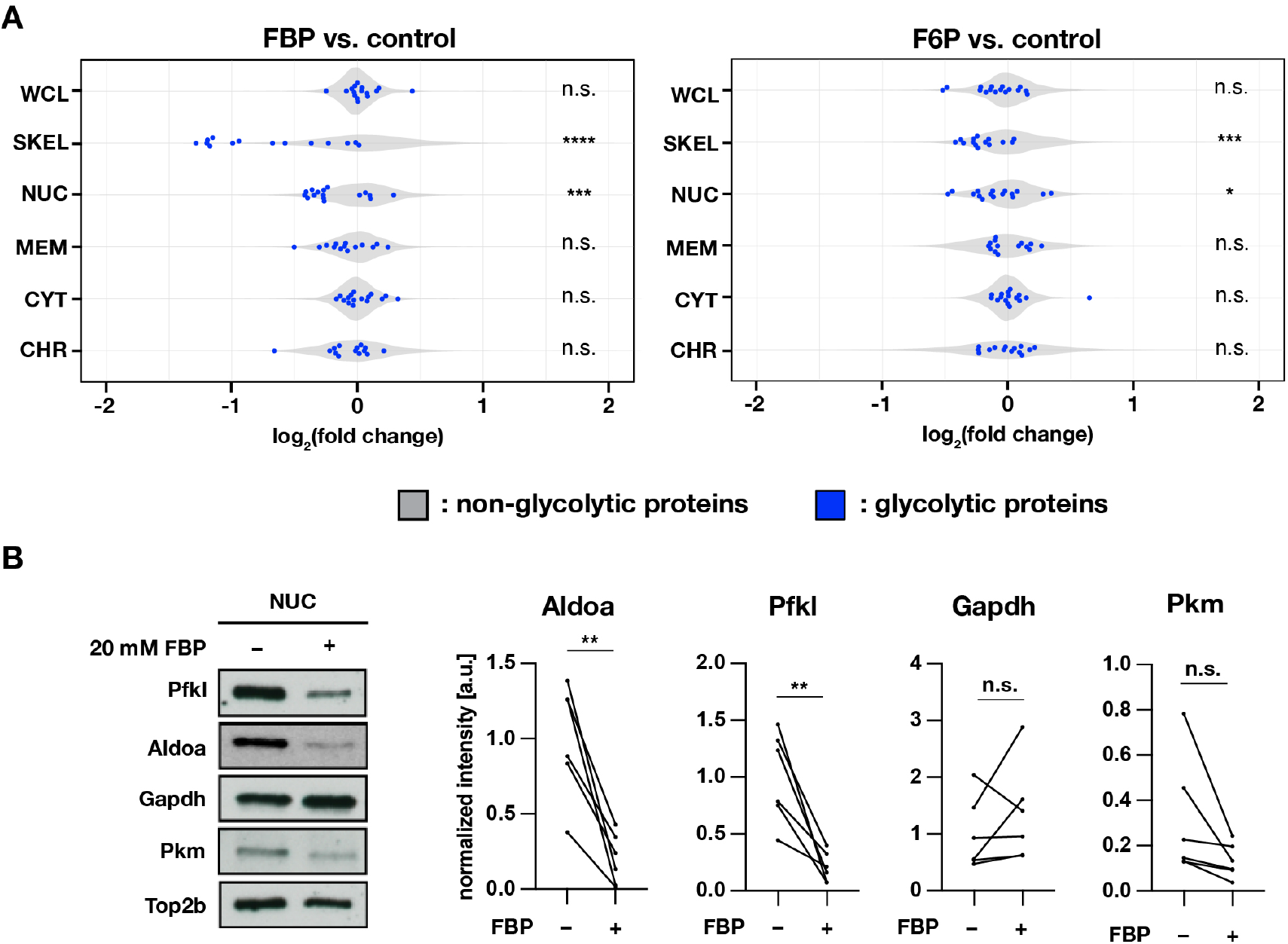
Subcellular localization of glycolytic enzymes are responsive to FBP treatment. **(A)** Effects of FBP treatment on subcellular localization of glycolytic enzymes. PSM explants were cultured for three hours in media containing 2.0 mM glucose and supplemented with 20 mM F6P or FBP. In addition to whole cell lysates (WCL), protein extracts were prepared from cytoplasmic (CYT), membrane (MEM), nuclear-soluble (NUC), chromatin-bound (CHR), and cytoskeletal (SKEL) fractions (n = 3 biological replicates). Abundance ratios (log_2_ (F6P/FBP-treated/control)) of glycolytic enzymes (in blue) were compared to those of non-glycolytic proteins (the rest, in gray) for statistical analysis (unpaired two-sample Wilcoxon test, \**p*-value < 0.05, \*\*\**p*-value < 0.001, \*\*\*\**p*-value < 0.0001, n.s., not significant). **(B)** Effects of FBP on the abundance of glycolytic enzymes in the nuclear soluble fraction. Subcellular protein fractionation was performed following one hour incubation of PSM explants in the media containing 0.5 mM glucose and supplemented with 20 mM FBP (n = 6 biological replicates; paired *t* -test, \*\**p*-value < 0.01, n.s., not significant).

We next asked whether subcellular localization of glycolytic enzymes is also altered upon cytoPfkfb3 overexpression, which we showed leads to an increase in glycolytic flux and FBP levels (Figure 3B, 3C). We hence performed sub-cellular proteome analysis of both control and *cytoPfkfb3* transgenic PSM explants, cultured for one hour in 10 mM glucose-containing medium. Due to the limited material obtained from transgenic embryos, proteins from nuclear-soluble, chromatin-bound, and cytoskeletal fractions were collected as a single, nuclear-cytoskeletal fraction. We found that cytoPfkfb3 overexpression altered the nuclear-cytoskeletal abundance of 12 proteins among 2813 detected proteins (adjusted *p*-value < 0.05 & |log_2_(fold change)| > 0.5) (Figure 7A-C). One of these proteins showing a pronounced depletion in the nuclear-cytoskeletal fraction in transgenic explants turned out to be the glycolytic enzyme Pfkl (Figure 7C). Using western blotting, we confirmed that Pfkl was depleted in the nuclear-cytoskeletal fraction in transgenic explants cultured at 10 mM glucose (Figure 7D). Importantly, under 2 mM glucose condition, nuclear-cytoskeletal Pfkl was not depleted in transgenic explants, suggesting that subcellular localization of Pfkl changes in a glucose-dose dependent manner. In addition, we found that, in *cytoPfkfb3* explants, the overall abundance of glycolytic machinery was decreased in the cytoplasmic and membrane fraction (Figure 7A-B). Combined, our results hence reveal that an alteration in glycolytic-flux/FBP levels, either by direct supplementation of metabolites or by genetic means using cytoPfkfb3 overexpression, changes the distribution of glycolytic enzymes in several subcellular compartments. While we have not been able to address the functional consequence of specific changes in subcellular localization, such as the nuclear depletion of Pfkl or Aldoa when glycolytic flux is increased, these results pave the way for future investigations on the mechanistic underpinning of how metabolic state is linked to cellular signaling and functions.

**Fig. 7.**
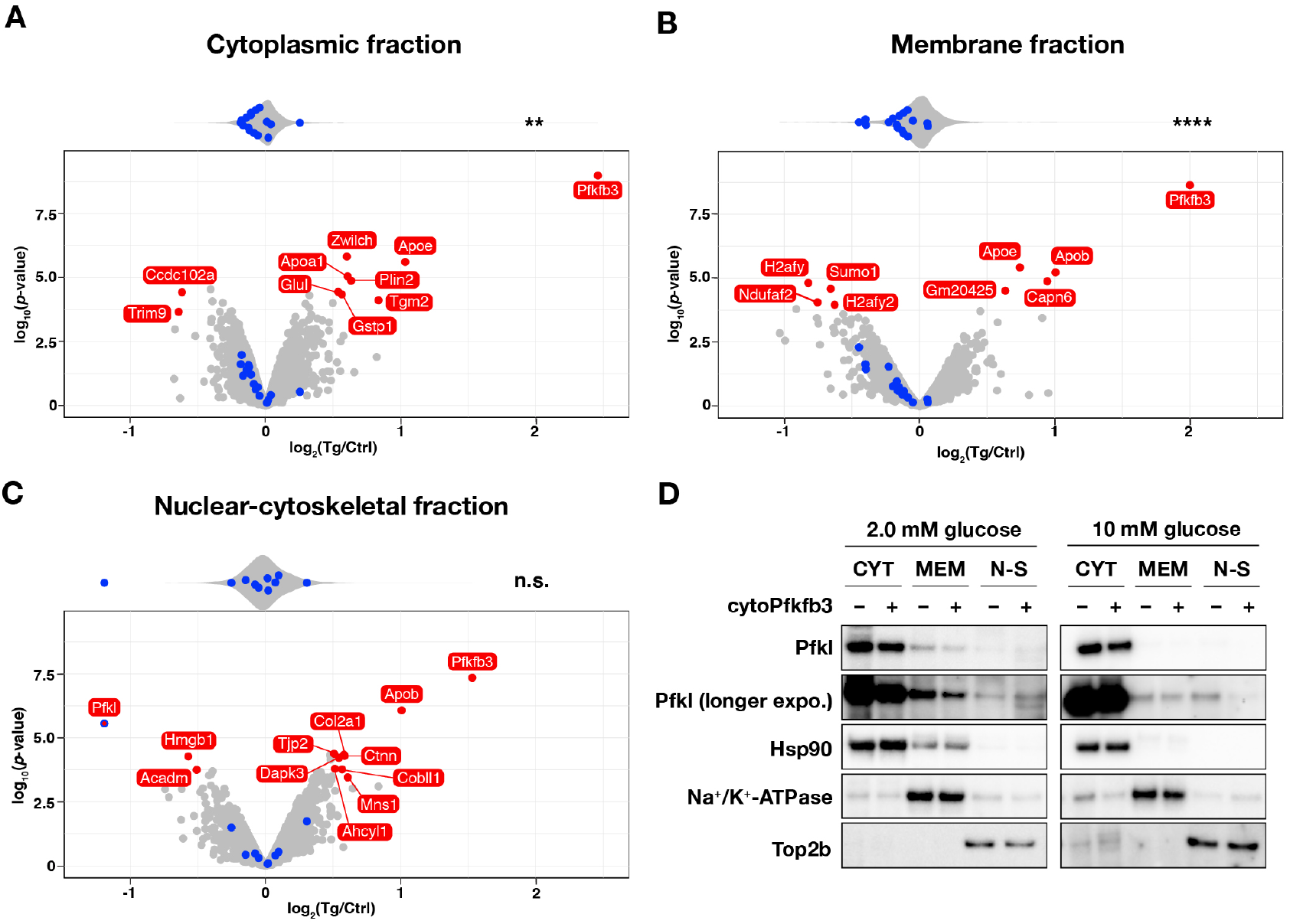
Subcellular localization of Pfkl responds to cytoPfkfb3 overexpression in a glucose-concentration dependent manner. **(A-C)** Effects of cytoPfkfb3 overexpression on subcellular protein localization assessed by mass spectrometry. Following one hour incubation of PSM explants in 10 mM glucose, protein extracts were prepared from cytoplasmic (A), membrane (B), and nuclear-cytoskeletal (C) fractions (n = 3 biological replicates). Proteins whose abundance showed significant changes (adjusted *p*-value < 0.05 & |log_2_ (fold change)| > 0.5) are marked red in the volcano plots. Top violin plots show distribution of abundance changes of glycolytic (blue) and non-glycolytic (gray) proteins. Statistical comparison as in Figure 6. \*\**p*-value < 0.01. Ctrl, control explants; Tg, *cytoPfkfb3* explants (crossed to the *Hprt-Cre* line). **(D)** Western-blot analysis of subcellular localization of Pfkl under different glucose conditions. Subcellular protein fractionation was performed following one hour incubation of PSM explants under 2.0 mM or 10 mM glucose (n = 3 biological replicates). CYT, cytoplasmic fraction; MEM, membrane fraction; N-S, nuclear-cytoskeletal fraction.

## Discussion

### Identifying FBP as a sentinel metabolite for glycolytic flux in developing mouse embryos

In this work, we investigated how glycolytic flux impacts mouse embryo mesoderm development, seeking to decipher the underlying mechanisms. First, we aimed to identify sentinel metabolites whose concentrations mirror glycolytic flux in mouse embryos (10, 42–44). The identification of sentinel metabolites is critical, as steady state metabolite levels are generally poor indicators of metabolic pathway activities (45). By investigating how steady state metabolite levels respond to an alteration in glycolytic flux upon glucose titration, we identified aspartate, FBP, and lactate as potential sentinel glycolytic metabolites whose steady state levels were either positively (*i*.*e*. FBP and lactate) or negatively (*i*.*e*. aspartate) correlated with extracellular glucose levels (Figure 1B). Similar changes were observed upon glycolytic activation by cytoPfkfb3 overexpression (Figure 3C). Remarkably, we found that FBP levels exhibit a strong linear correlation with a wide range of glucose concentrations, showing a 45-fold increase from 0.5 mM to 10 mM glucose conditions (Figure 1A). Previous studies suggested that the reversible reactions between FBP and PEP allow coupling of FBP to lower glycolytic flux (43), and importantly that feed-forward activation of pyruvate kinase by FBP enables the cell to establish a linear correlation between FBP and glycolytic flux over a wide range of FBP concentrations (43, 46). Such properties of lower glycolytic reactions may allow FBP to function as a generic sentinel metabolite for glycolytic flux in various biological contexts, from bacteria to mammalian cells (10, 33, 42, 43). This study extends such a finding of FBP as a glycolytic sentinel metabolite to *in vivo* mammalian embryos.

### FBP as a flux signaling metabolite connecting glycolytic-flux and PSM development

Interestingly, in addition to being a sentinel for glycolytic flux, FBP has been shown to carry signaling functions, hence relaying flux information to downstream effectors, such as transcription factors and signaling molecules (10, 42, 43). To test if such a flux-signaling function exists also in mouse embryos, we combined two complementary approaches, *i*.*e*. medium-supplementation of FBP (Figure 2) and, importantly, a genetic mouse model to increase glycolytic flux (Figure 3–5).

First, we revealed that high doses of FBP impaired mesoderm segmentation, disrupted the segmentation clock activity and led to downregulation of Wnt and Fgf target gene expression in the PSM (Figure 2). Using ^13^C-tracing experiments, we showed that exogenous FBP could be taken up by PSM cells (Figure S3C), an important control considering the debate regarding the permeability of this highly charged metabolite through the cell membrane (47). Interestingly, the effect of FBP appear most pronounced in the posterior, most undifferentiated PSM cells, while segmentation clock activity persists in the anterior PSM cells upon medium-supplementation of FBP (Figure 2A). This argues against a pleiotropic, toxic effect of FBP and suggests a more specific effect triggered by increased FBP levels. As a second, complementary approach to alter glycolytic flux and hence FBP levels, we aimed to increase glycolytic flux, and FBP levels, by acting on the activity of Pfk, the rate limiting glycolytic enzyme, in a genetic manner (Figure 3). To this end, we generated conditional transgenic mice which overexpress cytoPfkfb3 in a Cre-dependent manner. We showed that cytoPfkfb3 overexpression was indeed effective in increasing glycolytic flux in PSM explants, with a two-fold increase in secreted lactate (Figure 3B). Such a strong activation of glycolysis has been shown to be difficult to achieve by overexpression of single, wild-type glycolytic proteins in mammalian cell lines (33, 35). Of note, GC-MS analysis showed that cytoPfkfb3 overexpression was effective in increasing intracellular FBP levels (Figure 3C). Because the extent of glycolytic activation by cytoPfkfb3 was dependent on glucose concentration in the culture media (Figure 3B), we can titrate the effects of cytoPfkfb3 overexpression by increasing glucose. Therefore, the *cytoPfkfb3* transgenic mouse line that we generated is a powerful, genetic mouse model to study the function of glycolysis and, more importantly, that of a sentinel glycolytic metabolite FBP, in various biological contexts.

Functionally, overexpression of cytoPfkfb3 led to impairment of PSM segmentation at 10 mM or higher glucose concentrations, while wild-type PSM developed properly, at least qualitatively, at this glucose concentration (Figure 4, S1). The abnormal PSM development accompanied disruption of the segmentation clock activity and suppression of Wnt-target gene expression, while expression of FGF-target gene remained comparable to control. These phenotypes are reminiscent of our observation that intermediate levels (10 mM) of exogenous FBP suppressed mRNA expression of *Msgn* but not of *Dusp4* (Figure 2B). This data hence indicate that cytoPfkfb3 overexpression phenocopies the effect of the FBP-supplementation on PSM development.

Combined, these findings provide evidence that the sentinel glycolytic metabolite FBP exerts a signaling function in PSM development.

### The role of regulated flux at the level of PFK

One key finding we made is that PSM development and segmentation clock dynamics are particularly sensitive to changes in FBP levels that result from bypassing or impairing flux-regulation by Pfk. Hence, we found that supplementation of FBP, but not its precursor metabolite F6P, which is converted to FBP by Pfk, impaired mesoderm development and arrested segmentation clock oscillations. Along similar lines, cytoPfkfb3 overexpression, which interferes with allosteric regulation of Pfk, very effectively impaired PSM development and suppressed Wnt signaling target gene expression in a glucose-dose dependent manner. In contrast, the activation of glycolysis with intact flux-regulation by Pfk, such as the glucose titration experiment in control embryos, had relatively minor functional consequences on gene expression (Figure S3), segmentation and clock oscillation (Figure S1), while FBP levels were significantly altered (Figure 1).

We conclude from these results that flux-regulation by Pfk carries a critical role and offer several, non-exclusive interpretations. First, our findings might simply indicate that flux-regulation by Pfk plays a critical role in keeping FBP steady state levels within a critical range. In addition, these results are also compatible with a view in which flux-regulation by Pfk provides a critical dynamical control over glycolytic flux and FBP levels. While speculative at this point, our findings hence bear resemblance to previous classical findings showing the role of Pfk as a key regulator of glycolytic oscillations (48–50). Glycolytic oscillations and a function of metabolic rhythms have not been identified in embryonic development. Nevertheless, our results are compatible with the hypothesis that the regulation of FBP dynamics by Pfk could play a role during PSM patterning and segmentation clock oscillations. To probe for this possibility more directly, we are currently exploring strategies to investigate the presence of metabolic rhythms in embryonic cells using biosensors.

### Wnt signaling as a link between glycolytic-flux and the segmentation clock

We show that Wnt-target gene expression is responsive to glycolytic flux: while lowering glucose concentration correlated with an upregulation of Wnt target genes, the opposite effect was found when glucose concentration was increased (Figure S2). Consistently, we found that Wnt target gene expression showed a decrease in conditions of FBP supplementation and cytoPfkfb3 overexpression (Figure 2B, 5A). Previously, it was shown that Wnt signaling can promote glycolysis directly or indirectly (16, 51). Therefore, our findings suggest that in the PSM there is a negative-feedback regulation from glycolysis to Wnt signaling. Contrary to our findings, a previous study performed in cultured chick embryos has suggested that glycolysis regulates Wnt signaling positively, via the regulation of intracellular pH levels (30). While the reason for this discrepancy in how glycolysis impacts Wnt signaling is currently unknown, these combined findings do highlight that Wnt signaling is responsive to glycolytic-flux and hence a link between metabolism and PSM development. Given the central function of Wnt signaling in development, stem cells and disease, a future key interest will be to reveal its link to metabolism and in particular glycolytic flux in these different contexts. In addition, as FBP can be considered as a universal sentinel for glycolytic-flux in living organisms, it will be crucial to reveal the mechanisms of how cells integrate steady state FBP levels in these different contexts.

### Impact of altered glycolytic-flux and FBP levels on subcellular protein localization

As one mechanism by which FBP levels are integrated into cellular programs, we propose that FBP levels impact subcellular localization of proteins, some of which might function as FBP sensor molecules. Here, we revealed that several glycolytic enzymes including Aldoa and Pfkl are amongst those proteins altering their subcellular localization in response to FBP supplementation or cytoPfkfb3 overexpression (Figure 6, 7). Of note, we found that several glycolytic enzymes are, in the first place, localized in the nuclear soluble fraction (Figure S6C–E), raising the question about their functional role within the nuclear compartment. So far, glycolytic activity is thought to be restricted to the cytosol and hence nuclear localization of these enzymes raises the attractive hypothesis that their subcellular compartmentalization is linked to a non-metabolic, moonlighting function (52–56).

Additionally recent evidence in several biological systems highlights that subsets of metabolic reactions, for instance, from the mitochondrial TCA-cycle, take place also in the nucleus in order to maintain local supply of substrates for epigenetic modifications (57, 58). Thus, one possibility is that specific glycolytic reactions are taking place also in the nucleus, for instance to provide a local source of co-factors (*e*.*g*. NAD^+^) and/or substrates (*e*.*g*. acetyl-CoA, O-GlcNAc) for post-translational modifications of proteins. This emerging view of compartmentalized, local metabolic reactions as a way to regulate cellular functions has been recently supported by experimental evidence (41, 59–61).

While future studies will need to reveal if nuclear localization of glycolytic enzymes is linked to their moonlighting functions or metabolic compartmentalization, our finding that their subcellular localization is glycolytic flux-sensitive reveals a potentially general mechanism of how metabolic state is integrated into cellular programs. It is particularly interesting to find that Pfkl and Aldoa, enzymes that are known to directly bind to FBP, show a flux-sensitive subcellular localization (Figure 6, 7). This raises the possibility that FBP-protein interaction impacts, directly or indirectly, protein localization within a cell. Interestingly, a previous study has proposed that FBP is implicated in allosteric regulation of a multitude proteins involved in either metabolic as well as non-metabolic processes in *Escherichia coli* (62). Therefore, revealing the FBP allosterome and investigating the impact of allosteric interactions on protein localization and, more generally, on protein function, is of central importance and a key future objective (63). Excitingly, emerging techniques are now becoming available that enable interrogation of metabolite-protein interaction (62, 64) and we are currently exploring the possibility to decipher allosteromes in complex biological samples, such as in mouse embryos.

## Outlook

Using mouse embryo mesoderm development as a model system, our study identifies FBP as a sentinel, flux-signaling metabolite connecting glycolysis and developmental signaling pathways. Considering that cellular metabolism responds to various environmental cues, this integration of the metabolic state into cellular programs signifies an attractive mechanistic link of how environmental cues impact phenotype (65, 66). Interestingly, the role of FBP as a flux-signaling metabolite has been demonstrated in bacteria (3) and hence predates the origin of signaling pathways involved in multicellular organism development, such as the Wnt signaling pathway, which appeared only from the metazoa (67). From this perspective, it is of great interest to ask how metabolic flux-signaling has been integrated into signaling pathways involved in multicellular organism development in the course of evolution. Future studies addressing this fundamental question will shed a new light on potential roles of metabolism as regulators/modulators of organismal phenotype.

## Materials and Methods

Please refer to supplementary materials (Supplementary Note 1) for detailed materials and methods.

## Acknowledgements

We thank Theodore Alexandrov for helpful discussion and Jonathan Rodenfels and Joel I Perez-Perri for critical comments on the manuscript. We thank Jason Chesney for kindly sharing the Flag-Pfkfb3(K472A/K473A) plasmid, Yvonne Petersen for performing blastocyst injection of ESCs, Jana Kress for generating anti-Pfkl antibody, the nCounter core facility at the university of Heidelberg for expression analysis using NanoString technology, Bernd Klaus for helping with statistical analysis of the NanoString data and all the member of the Aulehla group for their support and helpful discussion. This work was supported by the EMBL Advanced Light Microscopy Facility (ALMF) and the Laboratory Animal Resources (LAR).

## Funding

The work was supported by: the European Molecular Biology Laboratory (K.P., M. B., A.A.); the EMBL International PhD Programme (M.T.S.; N.P.; E.K.); and the EMBL Interdisciplinary Postdoc (EI3POD) programme under H2020 Marie Skłodowska-Curie Actions COFUND (grant number 664726; H.M.; H.M.H.). H.M. was further supported by the Japan Society for the Promotion of Science (JSPS). H.M.H. was also supported by the Sigrid Juselius Foundation (Sigrid Juselius Stiftelse). This work further was supported by the German Research Foundation/DFG (project SFB 1324 – project number 331351713) to A.A.

## Author contributions

H.M. designed the project, performed the experiments, analyzed the data, wrote and revised the original manuscript. M.T.S. designed the project, performed the experiments, analyzed the data, and revised the manuscript. N.P. performed the NanoString analysis, analyzed the data, and commented on the manuscript. E.K. performed the GC-MS analysis, analyzed the data, and commented on the manuscript. H.M.H. performed the subcellular proteome analysis, analyzed the data, and commented on the manuscript. N.T. performed gene targeting of ESCs for generation of *Rosa26-stop*^*floxed*^*- cytoPfkfb3* mice. K.R.P. supervised the GC-MS analysis. M.B. supervised the subcellular proteome analysis. A.A. conceptualized and supervised the project, and wrote the manuscript.

## Competing financial interests

The authors declare that they have no conflict of interest.

## Supplementary Note 1: Materials and Methods

### *Ex vivo* culture of PSM explants

All animals were housed in the EMBL animal facility under veterinarians’ supervision and were treated following the guidelines of the European Commission, revised directive 2010/63/EU and AVMA guidelines 2007. The detection of a vaginal plug was designated as embryonic day (E) 0.5, and all experiments were conducted with E10.5 embryos. PSM explants with three intact somites were collected using micro scalpels (Feather Safety Razor, No. 715, 02.003.00.715) in DMEM/F12 (without glucose, pyruvate, glutamine, and phenol red; Cell Culture Technologies) supplemented with 0.5–25 mM glucose (Sigma-Aldrich, G8769), 2.0 mM glutamine (Sigma-Aldrich, G7513), 1.0% (w/v) BSA (Cohn fraction V; Equitech-Bio, BAC62), and 10 mM HEPES (Gibco, 15360-106). The explants were then washed with pre-equilibrated culture medium (DMEM/F12 supplemented with 0.5–25 mM glucose, 2.0 mM glutamine, and 1.0% (w/v) BSA) and were transferred to 8-well chamber slides (Lab-Tek, 155411) filled with 160 µl of the pre-equilibrated culture medium. When assessing the impacts of glycolytic intermediates on PSM development, culture medium supplemented with a glycolytic intermediate (*i*.*e*. fructose 1-phosphate (Sigma-Aldrich, F1127), fructose 6-phosphate (Sigma-Aldrich, F3637), fructose 1,6-bisphosphate (Santa Cruz, sc-221476), ^13^C_6_-fructose 1,6-bisphosphate (Cambridge Isotope laboratories, CLM-8962), 3-phosphoglycerate (Sigma-Aldrich, P8877)) was prepared with pre-equilibrated culture media right before dissection. Following *ex vivo* culture under 5% CO_2_, 60% O_2_ condition, the explants were washed with PBS and were fixed overnight with 4% (v/v) formaldehyde solution (Merck, 1040031000) at 4°C for further analyses.

### Generation of *Rosa26-stop*^*floxed*^*-cytoPfkfb3* mouse line

Flag-Pfkfb3(K472A/K473A) (hereafter termed as cytoPfkfb3) from (35) was amplified by PCR using the following primers: forward 5’-TAGGCCGGCCGCCACCATGGACTACAAGGACGACGACG-3’ and reverse 5’-TGGGCCGGCCGGAAATGGAATGGAACCGACAC-3’. The resulting amplicon was then cloned into the Rosa26 targeting vector Ai9 (68) using *F*seI restriction enzyme to generate the loxP-stop-loxP-Flag-cytoPfkfb3 construct. Conditional *cytoPfkfb3* transgenic mouse line was generated by standard gene targeting techniques using R1 embryonic stem cells. Briefly, chimeric mice were obtained by C57BL/6 blastocyst injection and then outbred to establish the line through germline transmission. *Rosa26-stop*^*floxed*^*-cytoPfkfb3* mouse line was maintained by crossing to CD1 mouse strain.

### Genotyping

The following mice used in this study were described previously and were genotyped using primers described in these references: *T-Cre* (36), *Hprt-Cre* (69), *LuVeLu* (23). The primers used for genotyping of *Rosa26-stop*^*floxed*^*-cytoPfkfb3* mice were as follows: forward 5’-GAGCTGCAGTGGAGTAGGCG-3’ and reverse 5’-CTCGACCATGGTAATAGCGA-3’ (predicted product size, 580 bp). The primers used for genotyping of *Rosa26-cytoPfkfb3* mice were as follows: forward 5’-GGCTTCTGGCGTGTGACCGG-3’ and reverse 5’-ACTCGGCTCTGCGTCAGTTC-3’ (predicted product size, 340 bp). For polymerase chain reaction (PCR), OneTaq 2X Master Mix with Standard Buffer was utilized (New England Biolabs).

### Time-lapse imaging of *LuVeLu* embryos

Imaging was performed as described before (70). In brief, samples were excited by 514 nm-wavelength argon laser or 960 nm-wavelength Ti:Sapphire laser (Chameleon-Ultra, Coherent) through 20× Plan-Apochromat objective (numerical aperture 0.8). In some experiments, samples were placed into agar wells (3% low Tm agarose, Biozyme, 840101) with 600 nm-width to restrain tissue movements during imaging. Image processing was done using the Fiji software (71).

### *In situ* hybridization

Fixed PSM explants were dehydrated with methanol and were stored at -20°C until use. Whole mount *in situ* hybridization was performed as described in (23).

### Gas chromatography-mass spectrometry (GC-MS) analysis

Wild-type and *cytoPfkfb3* transgenic PSM explants with no somite were cultured *ex vivo* for three hours under different glucose conditions, as described above. After washing twice with ice-cold PBS, the explants were snap frozen by liquid N_2_, and were stored at -80°C until use. Metabolites were extracted from the 25x explants by mechanically dissociating tissues by pipetting in 100 µl ice-cold methanol supplemented with ribitol (5.0 µg/mL) as an internal standard. For metabolite extraction from the conditioned medium, 20 µl of the medium was mixed with 40 µl of ice-cold methanol supplemented with ribitol. After incubation at 72°C for 15 min, one volume of ice-cold MilliQ water was added, followed by centrifugation at 14,000 rpm at 4°C for 10 mins. The supernatants were transferred to amber glass vials (Agilent, 5183-2073) and were dried by centrifugal evaporator EZ-2 Plus (SP Scientific) (30°C, Medium Boiling Point). The dried metabolite extracts were derivatized with 40 µL of 20 mg/mL methoxyamine hydrochloride (Alfa Aesar, 593-56-6) solution in pyridine (Sigma-Aldrich, 437611) for 90 min at 37°C, followed by addition of 80 µL N-methyl-trimethylsilyl-trifluoroacetamide (MSTFA) (Alfa Aesar, 24589-78-4) and 10-hour incubation at room temperature (72, 73). GC-MS analysis was performed using a Shimadzu TQ8040 GC-(triple quadrupole) MS system (Shimadzu Corp.) equipped with a 30m x 0.25 mm x 0.25 µm ZB-50 capillary column (7HG-G004-11; Phenomenex). One µL of the sample was injected in split mode (split ratio = 1:5) at 250°C using helium as a carrier gas with a flow rate of 1 mL/min. GC oven temperature was held at 100°C for 4 min followed by an increase to 320°C with a rate of 10 °C/min, and a final constant temperature period at 320°C for 11 min. The interface and the ion source were held at 280°C and 230°C, respectively. The detector was operated both in scanning mode (recording in the range of 50-600 m/z) as well as in MRM mode (for specified metabolites). For peak annotation, the GCMSsolution software (Shimadzu Corp.) was utilized. The metabolite identification was based on an in-house database with analytical standards utilized to define the retention time, the mass spectrum and marker ion fragments for all the quantified metabolites. The metabolite quantification was carried out by integrating the area under the curve of the MRM transition of each metabolite. The data were further normalized to the area under the curve of the MRM transition of ribitol.

### Liquid chromatography-mass spectrometry (LC-MS) analysis

After three-hour culture in the presence of 20 mM ^13^C_6_-FBP, PSM explants were washed with cold 154 mM ammonium acetate, snap frozen in liquid N_2_ and then dissociated in 0.5 mL ice-cold methanol/water (80:20, v/v) containing 0.20 µM of the internal standard lamivudine (Sigma-Aldrich, PHR1365). The resulting suspension was transferred to a reaction tube, mixed vigorously and centrifuged for 2 min at 16,000× g. Supernatants were transferred to a Strata® C18-E column (Phenomenex, 8B-S001-DAK) which were previously activated with 1 mL of CH_3_CN and equilibrated with 1 mL of MeOH/H_2_O (80:20, v/v). The eluate was dried in a vacuum concentrator. The dried metabolite extracts was dissolved in 50 µL 5 mM NH_4_OAc in CH_3_CN/H_2_O (75:25, v/v), and 3 µL of each sample was applied to an amide-HILIC (2.6 µm, 2.1 × 100 mm, Thermo Fisher, 16726-012105). Metabolites were separated at 30°C by LC using a DIONEX Ultimate 3000 UPLC system and the following solvents: solvent A consisting of 5 mM NH_4_OAc in CH_3_CN/H_2_O (5:95, v/v) and solvent B consisting of 5 mM NH_4_OAc in CH_3_CN/H_2_O (95:5, v/v). The LC gradient program was: 98% solvent B for 1 min, followed by a linear decrease to 40% solvent B within 5 min, then maintaining 40% solvent B for 13 min, then returning to 98% solvent B in 1 min and then maintaining 98% solvent B for 5 min for column equilibration before each injection. The flow rate was maintained at 350 µL/min. The eluent was directed to the hESI source of the Q Exactive mass spectrometer (QE-MS; Thermo Fisher Scientific) from 1.85 min to 18.0 min after sample injection. The scan range was set to 69.0 to 550 m/z with a resolution of 70,000 and polarity switching (negative and positive ionisation). Peaks corresponding to the calculated metabolites masses taken from an in-house metabolite library (MIM +/ H^+^ ± 2 mmU) were integrated using the El-MAVEN software (74).

### Extracellular lactate measurement

Condition medium was collected following 12-hour *ex vivo* culture of PSM explants, and was stored at -80C° until use. Fluorometric lactate measurements were performed with the Lactate Assay Kit (Biovision, K607) following manufacturer’s instructions with a slight modification. The reaction volume was reduced to 50 µl, and 0.5–1.0 µl of the conditioned medium was used for the analysis.

### Whole embryo roller-culture and TUNEL staining

Embryos were collected with the intact yolk sac at E8.5 in DMEM (1.0 g/L glucose, without glutamine and phenol red) (Gibco, 11880-028) supplemented with 2.0 mM glutamine, 10%(v/v) FCS, and 1%(v/v) penicillin/streptomycin (Gibco, 15140-122). The embryos were cultured for 24 hours using the roller bottle culture system in 50% rat serum/DMEM (supplemented with 2.0 mM glutamine and 1% (v/v) penicillin/streptomycin) under 8% CO_2_, 20% O_2_, and 72% N_2_ (flow rate, 20 mL/min) condition (38). Following the whole embryo culture, the embryos without the yolk sac and amniotic membrane were fixed with 4% formaldehyde overnight at 4°C. TUNEL staining was done with In Situ Cell Death Detection Kit (Roche, 12156792910) following manufacturer’s instructions, followed by DAPI (0.5 µg/mL) staining. Images were acquired with a LSM780 laser-scanning microscope (Zeiss) using 10× EC Plan-Neofluar objective lens (numerical aperture 0.3).

### Subcellular proteome analysis by mass spectrometry

#### Effect of FBP and F6P on subcellular localization

Treated mouse PSMs were washed twice with ice-cold PBS and subjected to subcellular protein extraction using a Subcellular Protein Fractionation for Cultured Cells kit (Thermo Fisher Scientific, #78840). 8–11x PSMs were used for each condition in each replicate. PSMs were dissociated in 10 ul of CEB buffer per PSM by pipetting, after which 10 ul (*i*.*e*. 1× PSM worth) of uncleared lysate was taken as the whole-cell lysate (WCL) sample. The rest of the extraction was carried out following manufacturer’s instructions using buffer amounts scaled according to the number of PSMs in the sample. The resulting fractions were stored at -80°C before further processing. Subsequently, CYT and MEM fractions were reduced in volume to ∼50 ul in a speedvac, and each subcellular protein fraction was denatured with 1% SDS at 95°C for 5 minutes, after which residual nucleic acids were degraded with benzonase (EMD Millipore, #71206-25KUN; final concentration 0.1-1 U/µl) for 45 minutes at 37°C and 300 rpm until samples were no longer viscous.

#### Subcellular proteomics of *cytoPfkfb3* transgenic embryos

Transgenic (Tg) and control (Ctrl) PSM explants were subjected to subcellular protein extraction as above with the following exceptions: After extraction of the MEM fraction, protein from the remaining pellet (constituting the nuclear and cytoskeletal fractions) was extracted with the NEB buffer (with micrococcal nuclease) plus 1× SDS lysis buffer [50 mM HEPES-NaOH (pH 8.5), 1% SDS, 1x cOmplete protease inhibitor cocktail (Roche, 11873580001)] and used directly for benzonase treatment as above.

#### Sample preparation and LC-MS/MS

All samples were prepared for MS using a modified SP3 protocol (75). Briefly, protein samples were precipitated onto Sera-Mag SpeedBeads (GE Healthcare, #45152105050250 and #65152105050250) in the presence of 50% ethanol and 2.5% formic acid (FA) for 15 min at room temperature, followed by four washes with 70% ethanol on magnets. Proteins were digested on beads with trypsin and Lys-C (5 ng/µl final concentration each) in 90 mM HEPES (pH 8.5), 5 mM chloroacetic acid and 1.25 mM TCEP overnight at room temperature shaking at 500 rpm. Peptides were eluted on magnets using 2% DMSO and dried in a speedvac. Dry peptides were reconstituted in 10 µl water and labelled by adding 4 µl TMT label (20 µg/µl in acetonitrile (ACN)) (TMT10plex, Thermo Fisher Scientific #90110, comparison of FBP and F6P treatment; or TMTsixplex, #1861431, comparison of TG to Ctrl) and incubating for one hour at room temperature. Samples were multiplexed as follows: for comparison of FBP and F6P treatment all conditions (FBP, F6P, untreated) of each biological replicate were run in two separate TMT sets: one set including WCL, CYT and NUC fractions, and the other MEM, CHR and SKEL, for a total of six TMT sets for the experiment. For the comparison of Tg and Ctrl, both conditions (Tg, Ctrl) and all replicates were run in a single TMTsixplex experiment for each subcellular fraction (nuclear-cytoskeletal, cytoplasmic, membrane). Labeling was quenched with hydroxylamine (1.1% final concentration), and samples were dried in a speedvac. Each sample was then resuspended using 100 ul LC-MS H_2_O, and 10% of each sample was taken, pooled to make a full TMT set, and desalted on an OASIS HLB µElution plate (Waters 186001828BA); washing twice with 0.05% FA, eluting with 80% ACN, 0.05% FA, and drying in a speedvac. The resulting sample was run on a 60 min LC-MS/MS gradient (see details below) to estimate relative amounts of protein in each channel. A second TMT set was then pooled using equalized amounts based on the median intensity of each channel from the first run to create an analytical TMT set with approximately equal labelled input protein in each channel. The analytical TMT set peptides were desalted using OASIS as described above and dried in a speedvac. Dried peptides were taken up in 20 mM ammonium formate (pH 10) and prefractionated offline into six (comparison of FBP, F6P to untreated) or 12 (comparison of Tg to Ctrl) fractions on an Ultimate 3000 (Dionex) HPLC using high-pH reversed-phase chromatography (running buffer A: 20 mM ammonium formate pH 10; elution buffer B: ACN) on an X-bridge column (2.1 × 10 mm, C18, 3.5 µm, Waters). Prefractionated peptides were vacuum dried.

For LC-MS/MS analysis, peptides were reconstituted in 0.1% FA, 4% ACN and analyzed by nanoLC-MS/MS on an Ultimate 3000 RSLC (Thermo Fisher Scientific) connected to a Fusion Lumos Tribrid (Thermo Fisher Scientific) mass spectrometer, using an Acclaim C18 PepMap 100 trapping cartridge (5 µm, 300 µm i.d. x 5 mm, 100 Å) (Thermo Fisher Scientific) and a nanoEase M/Z HSS C18 T3 (100Å, 1.8 µm, 75 µm x 250 mm) analytical column (Waters). Solvent A: aqueous 0.1% FA; Solvent B: 0.1% FA in ACN (all LC-MS grade solvents are from Thermo Fisher Scientific). Peptides were loaded on the trapping cartridge using solvent A for 3 min with a flow of 30 µl/min. Peptides were separated on the analytical column with a constant flow of 0.3 µl/min applying a 120 min gradient of 2–40% of solvent B in solvent A. Peptides were directly analyzed in positive ion mode with a spray voltage of 2.2 kV and a ion transfer tube temperature of 275 °C. Full scan MS spectra with a mass range of 375–1500 m/z were acquired on the orbitrap using a resolution of 120,000 with a maximum injection time of 50 ms. Data-dependent acquisition was used with a maximum cycle time of 3 s. Precursors were isolated on the quadrupole with an intensity threshold of 2e5, charge state filter of 2-7, isolation window of 0.7 m/z. Precursors were fragmented using HCD at 38% collision energy, and MS/MS spectra were acquired on the orbitrap with a resolution of 30 000, maximum injection time of 54 ms, normalized AGC target of 200%, with a dynamic exclusion window of 60 s.

#### Data analysis

The mass spectrometry proteomics data have been deposited to the ProteomeXchange Consortium via the PRIDE (76) partner repository with the dataset identifier PXD029988. Mass spectrometry raw files were processed using IsobarQuant (77) and peptide and protein identification was obtained with Mascot 2.5.1 (Matrix Science) using a reference mouse proteome (uniprot Proteome ID: UP000000589, downloaded 14.5.2016) modified to include known common contaminants and reversed protein sequences. Mascot search parameters were: trypsin; max. 2 missed cleavages; peptide tolerance 10 ppm; MS/MS tolerance 0.02 Da; fixed modifications: Carbamidomethyl (C), TMT10plex (K); variable modifications: Acetyl (Protein N-term), Oxidation (M), TMT10plex (N-term).

IsobarQuant output data was analyzed on a protein level in R using in-house data analysis pipelines. In brief, protein data was filtered to remove contaminants, proteins with less than 2 unique quantified peptide matches as well as proteins, which were only detected in a single replicate. Subsequently, protein reporter signal sums were normalized within each TMT set using the vsn package (78). Significantly changing proteins between the treated and untreated sample were identified by applying a limma analysis (79) on the vsn-corrected values. Replicates were treated as covariates in the limma analysis for the comparison of FBP to F6P, as biological replicates were run as separate TMT sets. Multiple-testing adjustment of *p* values was done using the Benjamini-Hochberg method.

### Western blot analysis

Primaries antibodies used in the study are as follows: anti-Aldolase A (Proteintech, 11217-1-AP, 1:5,000), anti-Tpi (Acris, AP16324PU-N, 1:5,000), anti-Gapdh (Millipore, MAB374, 1:5,000), anti-Pkm1/2 (Cell signaling, 3190, 1:5000), anti-Histone H2B (Millipore, 07-371, 1:10,000), anti-beta-Tubulin (Millipore, 05-661, 1:10,000), anti-Hsp90 (Cell signaling, 4874, 1:1,000). Mouse monoclonal antibody against Pfkl was generated by EMBL Monoclonal Antibody Core Facility using full-length Pfkl as an antigen. For protein expression and purification, full-length Pfkl transcript was amplifed by reverse transcription (RT)–PCR using mouse embryo total RNA as a template and cloned into pET28M-SUMO3 vector (EMBL Protein Expression and Purification Core Facility) using *A*geI and *N*otI restriction enzymes. Following primers were used for RT-PCR: forward 5’-TCATCTACCGGTGGAATGGCTACCGTGGACCTGGAGA-3’ and reverse 5’-TCATCTGCGGCCGCTCAGAAACCCTTGTCTATGCTCAAGGT-3’.

### Gene expression analysis by NanoString nCounter Analysis System

A custom probe set was designed to include 237 genes involved in glucose metabolism, Notch-, Wnt-, and FGF-signaling pathways. In addition, six positive controls, eight negative controls and housekeeping genes for normalisation (housekeeping genes used: *Cltc, Gusb, Hprt1* and *Tubb5*) were included in the probe set. Following three-hour culture with the specified glucose concentration, the PSM explants were further dissected immediately posterior to the neural tube to isolate the posterior PSM. Five posterior PSM samples were pooled per replicate and snap frozen by liquid N_2_. Total RNA was isolated using TRIzol reagent (Invitrogen) according to manufacturer’s instructions and concentrated using RNA Clean & Concentrator-5 Kit (Zymo research). RNA was hybridized to the probes at 65 °C, samples were inserted into the nCounter Prep Station for 3 hours, the sample cartridge was transferred to the nCounter Digital Analyzer, and counts were determined for each target molecule. Counts were analysed using nSolver Analysis Software Version 4.0, and sequentially subjected to background correction, positive control (quality control) and normalisation to housekeeping genes.

### Statistical analysis

Statistical analysis was performed with GraphPad PRISM 9 software. For the metabolome data, statistical analysis was performed with the Statistical Analysis for Microarray (SAM) package (80) using R. For Pearson correlation analysis, numpy (81), pandas (82), and scipy (83) libraries were used. For data visualization, matplotlib (84) library was utilized.

**Fig. S1.**
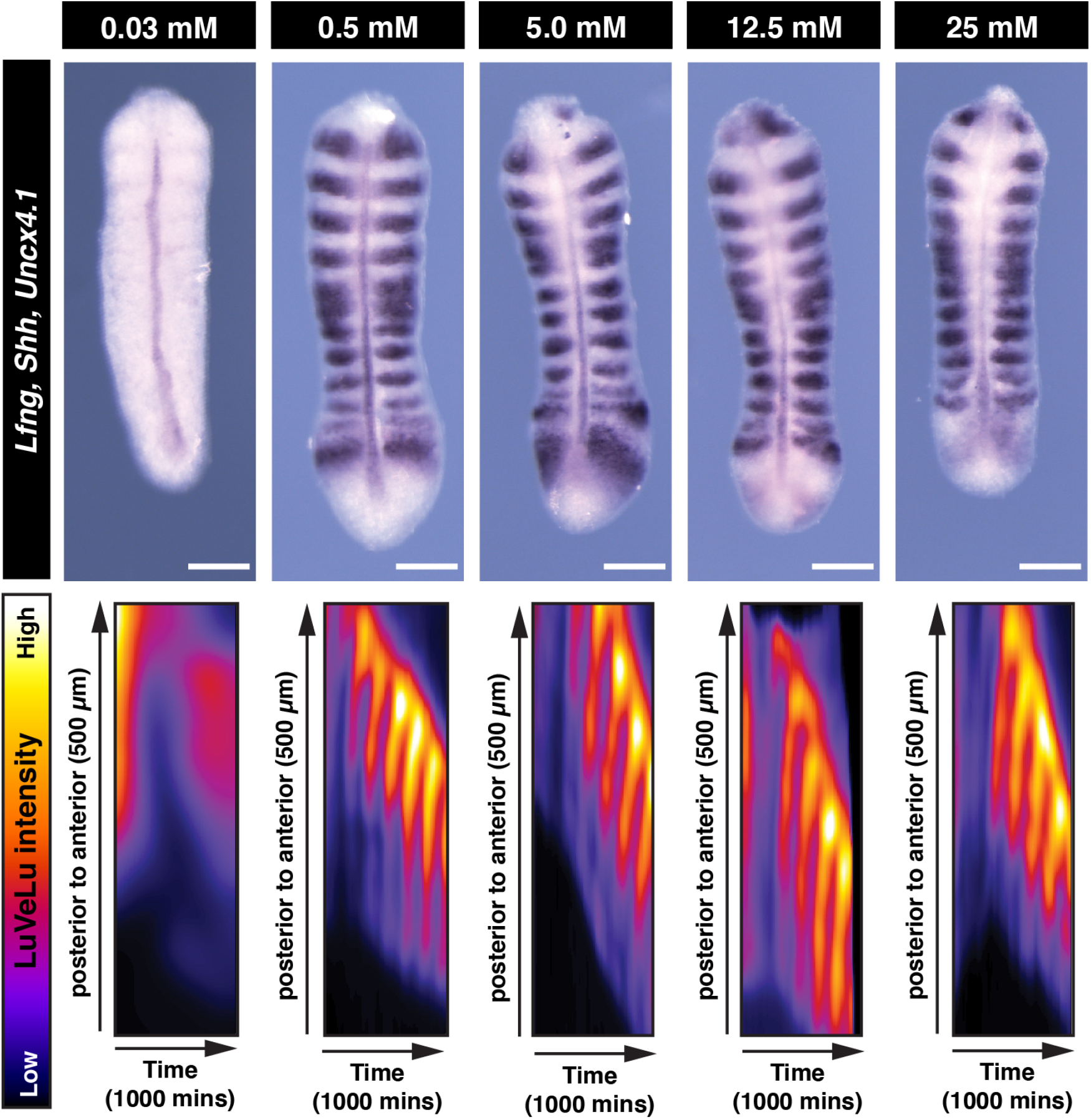
PSM patterning is tolerated within a wide range of glucose concentrations. Whole mount *in situ* hybridization analysis for *Lfng, Shh*, and *Uncx4*.*1* gene expressions in the PSM. PSM explants were incubated for 13 hours *ex vivo* in different glucose conditions. Kymographs show ongoing oscillatory dynamics of the Notch signaling activity reporter LuVeLu in the PSM in all conditions but 0.03 mM glucose condition. Scale bar, 100 µm.

**Fig. S2.**
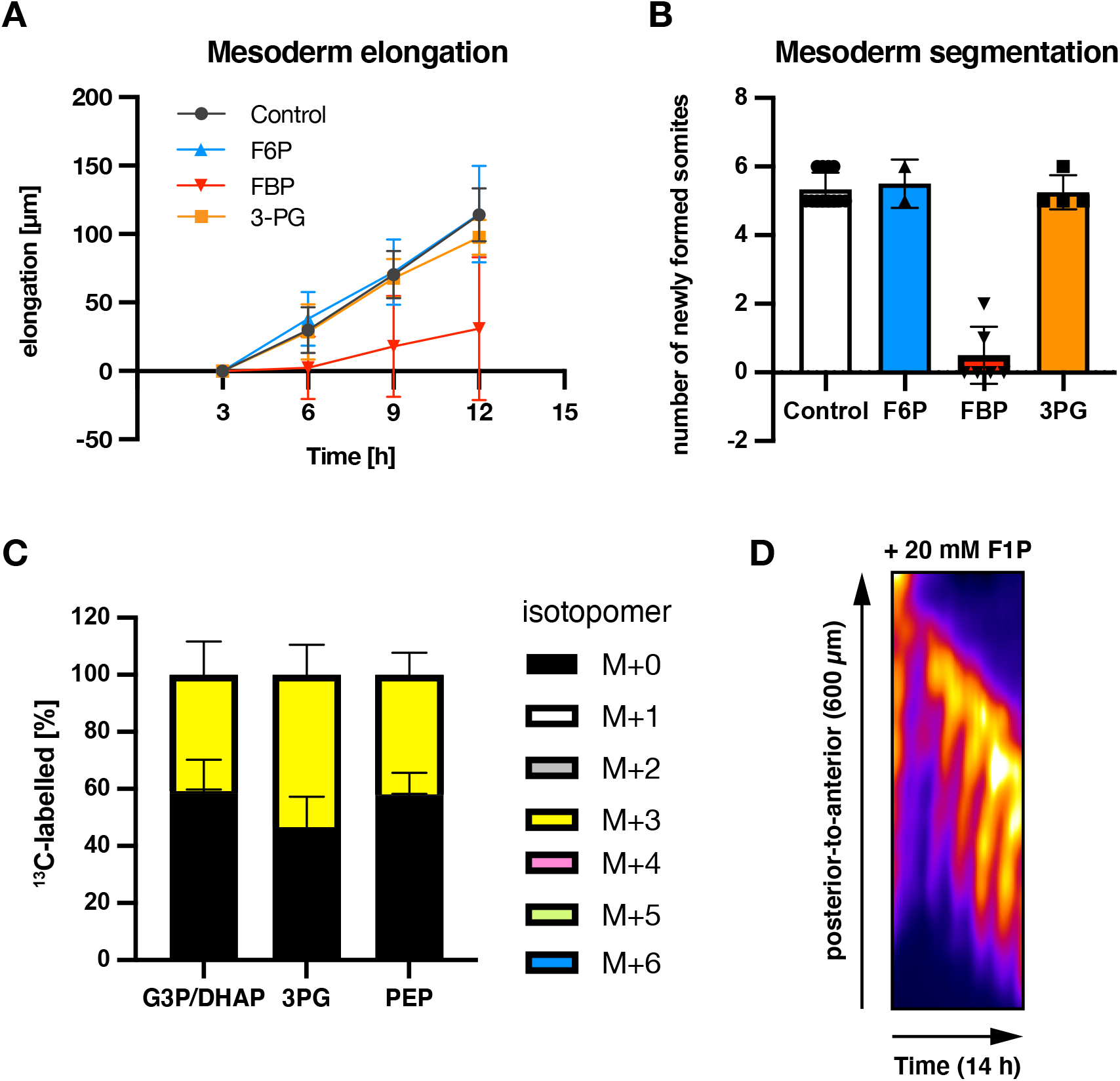
Effects of medium-supplementation of glycolytic intermediates on mesoderm elongation and segmentation. **(A)** Elongation of PSM explants during *ex vivo* culture. The explants were cultured in the medium containing 0.5 mM glucose and supplemented with 20 mM of F6P/FBP/3PG. The length of explants at three hour-incubation was used as the reference. **(B)** The number of newly formed somites during 12-hour *ex vivo* incubation. **(C)** ^13^C-tracing experiments with fully ^13^C-labelled FBP (^13^C_6_-FBP). The PSM explants were cultured for three hours in the medium containing 2.0 mM of glucose and supplemented with 20 mM ^13^C_6_-FBP. **(D)** Kymographs showing dynamics of the Notch signaling activity reporter LuVeLu in the PSM. Explants were cultured in medium containing 0.5 mM glucose and supplemented with 20 mM fructose 1-phosphate (F1P).

**Fig. S3.**
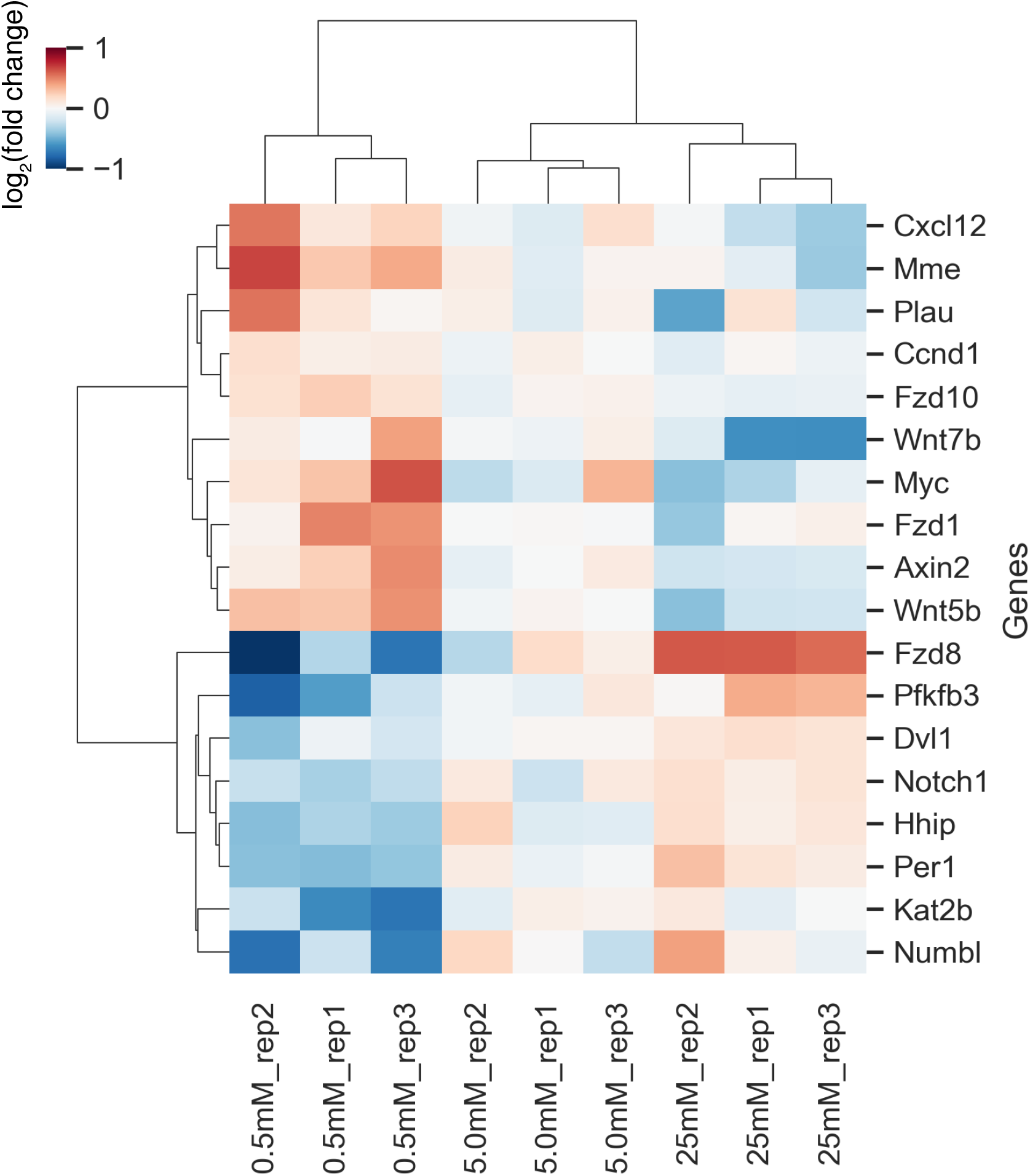
Modulation of Wnt-target gene expression upon glucose titration within PSM cells. Hierarchical clustering heatmap of genes whose expression levels showed linear correlation with extracellular glucose concentrations (linear regression analysis; adjusted *p*-value < 0.1). Following three-hour culture at varying concentrations of glucose, expressions of 237 genes were analyzed in the posterior PSM using the NanoString nCounter Analysis System. These genes included ones involved in glucose metabolism, Notch-, Wnt-, and FGF-signaling pathways. Fold changes were calculated using 5.0 mM glucose condition as the reference. Hierarchical clustering was performed using Ward’s method with Euclidean distance.

**Fig. S4.**
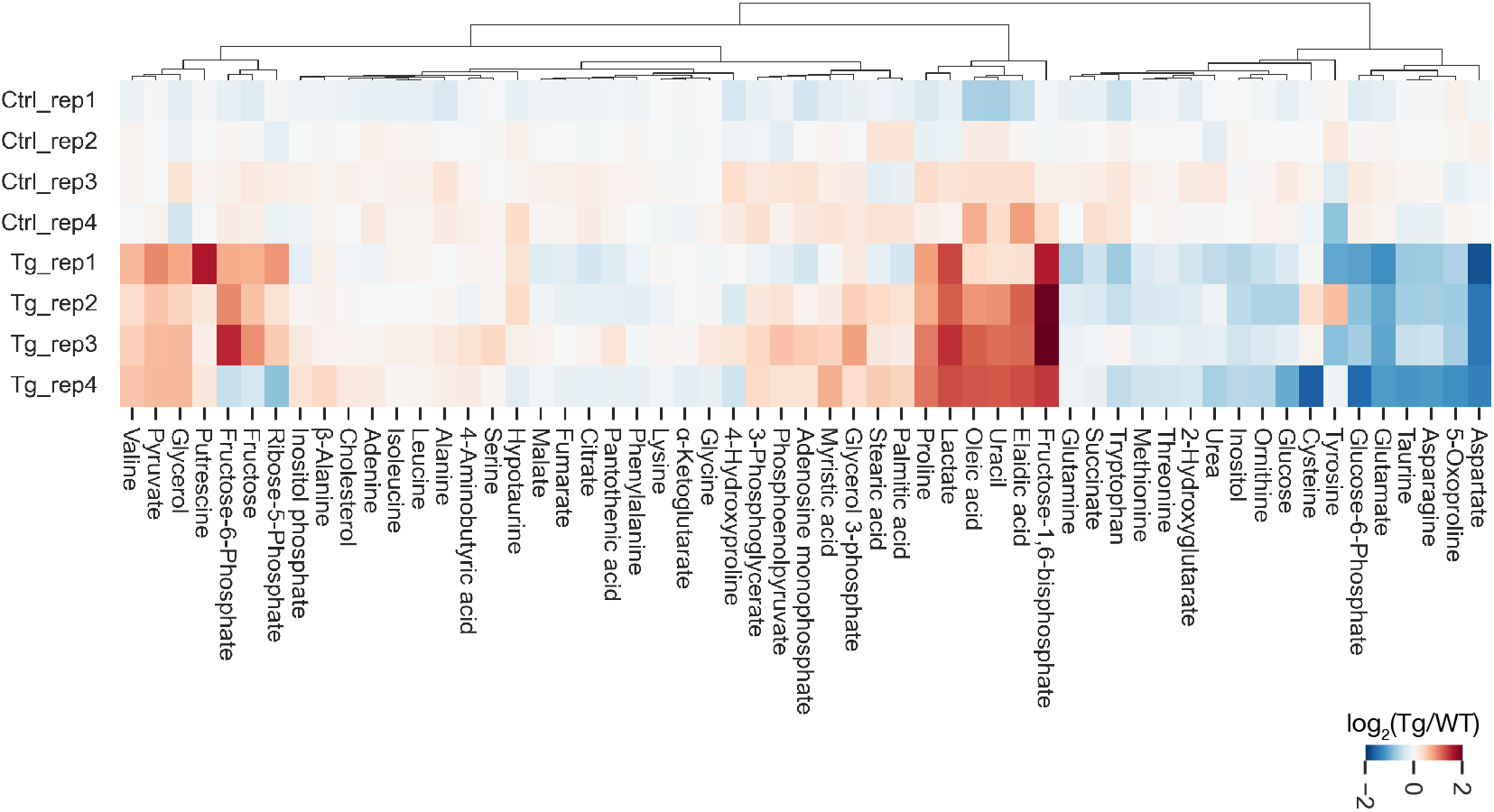
Steady state measurements of metabolites within *cytoPfkfb3* and control PSM explants by GC-MS. Hierarchical clustering heatmap of metabolites detected in PSM explants. Metabolomics analysis was performed following three-hour culture of PSM explants at 10 mM glucose cocentration (n = 4 biological replicates for each condition). Hierarchical clustering was performed using Ward’s method with Euclidean distance. Ctrl, control explants; Tg, *cytoPfkfb3* explants (crossed to *Hprt-Cre* line).

**Fig. S5.**
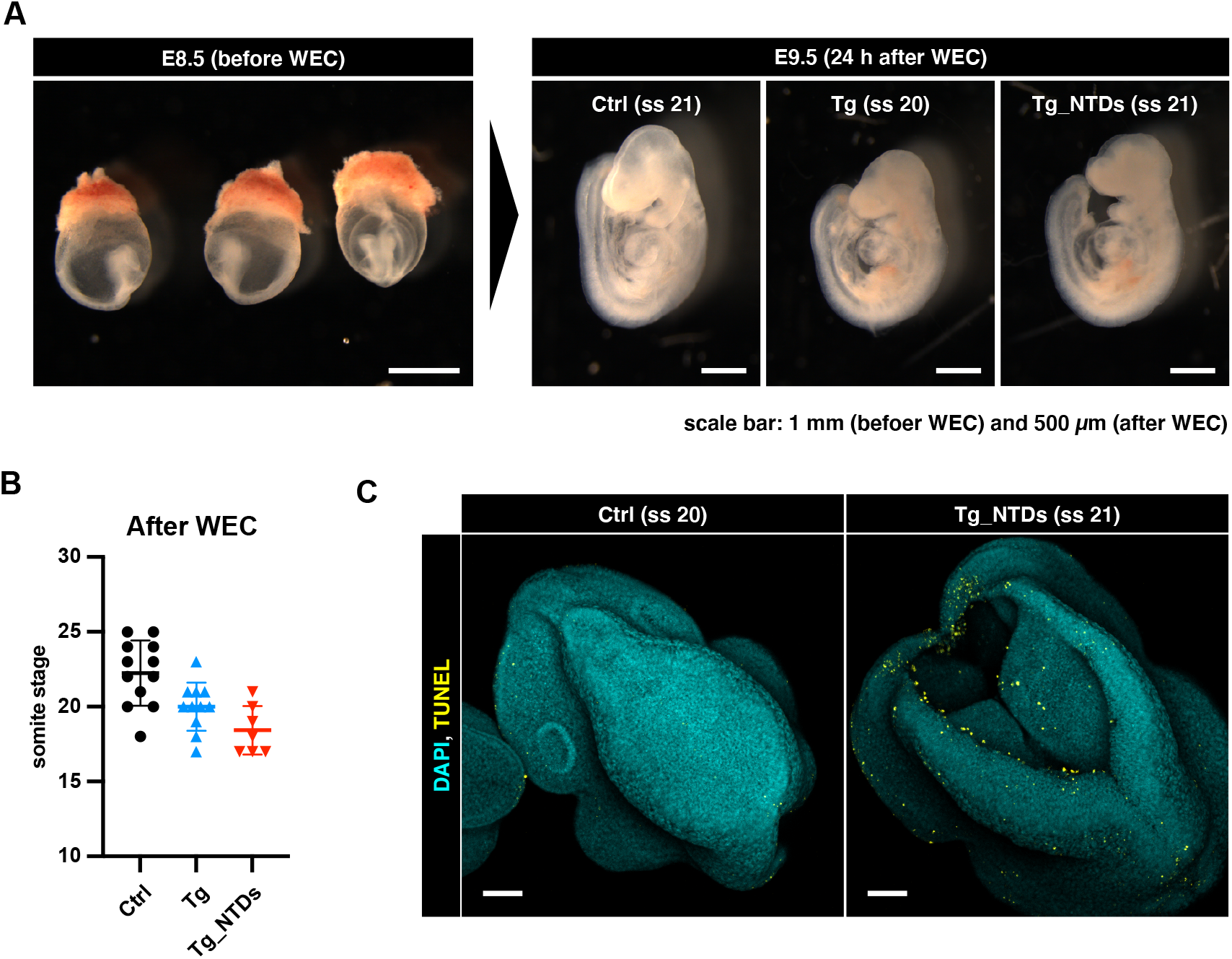
*cytoPfkfb3* transgenic embryos show defects in neural tube closure under normoglycemic conditions. **(A)** E8.5 embryos with the intact yolk sac were cultured for 24 hours at normoglycemic conditions using the whole embryo roller-culture (WEC) system. **(B)** Total number of somites after 24-hour of WEC. Data are represented as mean ± s.d. Ctrl, control embryos; Tg, *cytoPfkfb3* embryos (crossed to *Hprt-Cre* line); Tg_NTDs, *cytoPfkfb3* embryos with neural tube closure defects. **(C)** DAPI and TUNEL staining of the embryos following WEC. Midbrain region of the embryos is shown. Scale bar, 100 µm (ss, somite stage).

**Fig. S6.**
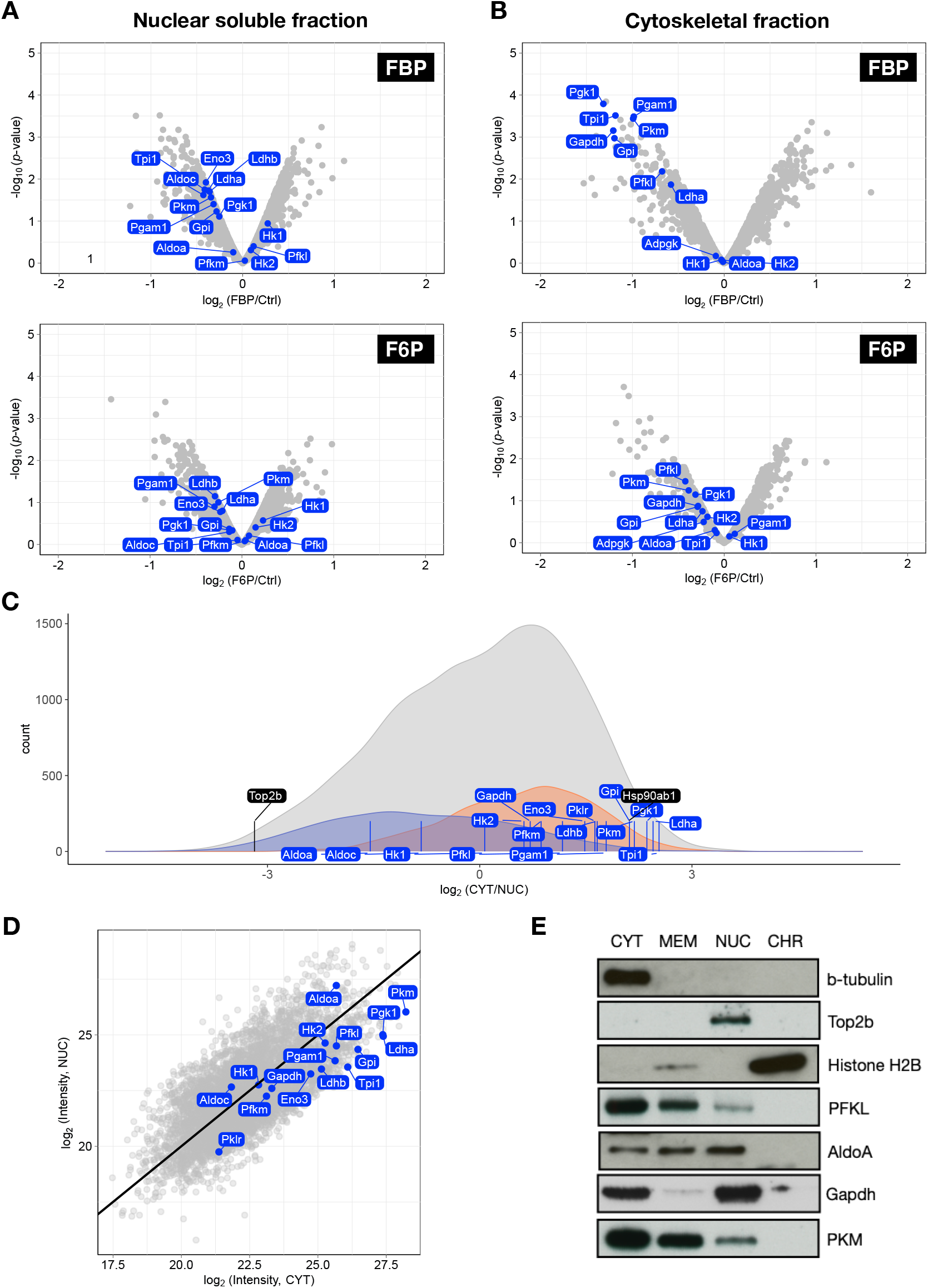
Proteome analysis of subcellular protein localization in the PSM. **(A, B)** Volcano plots showing effects of FBP-or F6P-treatment on abundance of proteins in nuclear-soluble (A) or cytoskeletal fractions (B). Subcellular protein fractionation was performed following three-hour culture of PSM explants in the medium containing 2.0 mM glucose and supplemented with 20 mM FBP or F6P. Glycolytic proteins are highlighted in blue. **(C)** Density plot showing the number of proteins that are annotated to nuclear (shown in blue) or cytoplasmic (shown in orange) compartments (based on GO term). PSM explants cultured in the control medium containing 2.0 mM gluocse were used for the analysis. All the detected-proteins are shown in gray. Glycolytic proteins are highlihgted by blue lines, and marker proteins for the nuclear-soluble (*i*.*e*. Top2b) and cytoplasmic (*i*.*e*. Hsp90ab1) fractions are highlihgted by black lines. **(D)** Abundance ratio of glycolytic proteins (marked by blue) between nuclear-soluble (NUC) and cytoplasmic (CYT) fractions. **(E)** Western blot analysis of glycolytic proteins following subcellular protein fractionation of the PSM explants cultured in the control medium containing 0.5 mM glucose.

## References

1. A. Efeyan, W. C. Comb, and D. M. Sabatini. Nutrient-sensing mechanisms and pathways. Nature, 517(7534):302–10, 2015. ISSN 1476-4687 (Electronic) 0028-0836 (Linking). doi: 10.1038/nature14190.

2. Jr. Kaelin, W. G. and P. J. Ratcliffe. Oxygen sensing by metazoans: the central role of the hif hydroxylase pathway. Mol Cell, 30(4):393–402, 2008. ISSN 1097-4164 (Electronic) 1097-2765 (Linking). doi: 10.1016/j.molcel.2008.04.009.

3. A. Litsios, A. D. Ortega, E. C. Wit, and M. Heinemann. Metabolic-flux dependent regulation of microbial physiology. Curr Opin Microbiol, 42:71–78, 2018. ISSN 1879-0364 (Electronic) 1369-5274 (Linking). doi: 10.1016/j.mib.2017.10.029.

4. Y. P. Wang and Q. Y. Lei. Metabolite sensing and signaling in cell metabolism. Signal Transduct Target Ther, 3:30, 2018. ISSN 2059-3635 (Electronic) 2059-3635 (Linking). doi: 10.1038/s41392-018-0024-7.

5. R. A. Saxton and D. M. Sabatini. mtor signaling in growth, metabolism, and disease. Cell, 168(6):960–976, 2017. ISSN 1097-4172 (Electronic) 0092-8674 (Linking). doi: 10.1016/j.cell.2017.02.004.

6. A. Gonzalez, M. N. Hall, S. C. Lin, and D. G. Hardie. Ampk and tor: The yin and yang of cellular nutrient sensing and growth control. Cell Metab, 31(3):472–492, 2020. ISSN 1932-7420 (Electronic) 1550-4131 (Linking). doi: 10.1016/j.cmet.2020.01.015.

7. S. L. Campbell and K. E. Wellen. Metabolic signaling to the nucleus in cancer. Mol Cell, 71(3):398–408, 2018. ISSN 1097-4164 (Electronic) 1097-2765 (Linking). doi: 10.1016/j.molcel.2018.07.015.

8. M. A. Reid, Z. Dai, and J. W. Locasale. The impact of cellular metabolism on chromatin dynamics and epigenetics. Nat Cell Biol, 19(11):1298–1306, 2017. ISSN 1476-4679 (Electronic) 1465-7392 (Linking). doi: 10.1038/ncb3629.

9. H. Miyazawa and A. Aulehla. Revisiting the role of metabolism during development. Development, 145(19), 2018. ISSN 1477-9129 (Electronic) 0950-1991 (Linking). doi: 10.1242/dev.131110.

10. K. Peeters, F. Van Leemputte, B. Fischer, B. M. Bonini, H. Quezada, M. Tsytlonok, D. Haesen, W. Vanthienen, N. Bernardes, C. B. Gonzalez-Blas, V. Janssens, P. Tompa, W. Versees, and J. M. Thevelein. Fructose-1,6-bisphosphate couples glycolytic flux to activation of ras. Nat Commun, 8(1):922, 2017. ISSN 2041-1723 (Electronic) 2041-1723 (Linking). doi: 10.1038/s41467-017-01019-z.

11. M. T. Snaebjornsson and A. Schulze. Non-canonical functions of enzymes facilitate crosstalk between cell metabolic and regulatory pathways. Exp Mol Med, 50(4):1–16, 2018. ISSN 2092-6413 (Electronic) 1226-3613 (Linking). doi: 10.1038/s12276-018-0065-6.

12. A. E. Boukouris, S. D. Zervopoulos, and E. D. Michelakis. Metabolic enzymes moonlighting in the nucleus: Metabolic regulation of gene transcription. Trends Biochem Sci, 41(8):712– 730, 2016. ISSN 0968-0004 (Print) 0968-0004 (Linking). doi: 10.1016/j.tibs.2016.05.013.

13. C. H. Chang, J. D. Curtis, Jr. Maggi, L. B., B. Faubert, A. V. Villarino, D. O’Sullivan, S. C. Huang, G. J. van der Windt, J. Blagih, J. Qiu, J. D. Weber, E. J. Pearce, R. G. Jones, and E. L. Pearce. Posttranscriptional control of t cell effector function by aerobic glycolysis. Cell, 153(6):1239–51, 2013. ISSN 1097-4172 (Electronic) 0092-8674 (Linking). doi: 10.1016/j.cell.2013.05.016.

14. Jr. Spratt, N. T. Nutritional requirement of the early chick embryo. iii. the metabolic basis of the morphogenesis and differentiation as revealed by the use of inhibitors. Biol Bull, 99(1): 120–35, 1950. ISSN 0006-3185 (Print) 0006-3185 (Linking). doi: 10.2307/1538756.

15. V. Bulusu, N. Prior, M. T. Snaebjornsson, A. Kuehne, K. F. Sonnen, J. Kress, F. Stein, C. Schultz, U. Sauer, and A. Aulehla. Spatiotemporal analysis of a glycolytic activity gradient linked to mouse embryo mesoderm development. Dev Cell, 40(4):331–341 e4, 2017. ISSN 1878-1551 (Electronic) 1534-5807 (Linking). doi: 10.1016/j.devcel.2017.01.015.

16. M. Oginuma, P. Moncuquet, F. Xiong, E. Karoly, J. Chal, K. Guevorkian, and O. Pourquie. A gradient of glycolytic activity coordinates fgf and wnt signaling during elongation of the body axis in amniote embryos. Dev Cell, 40(4):342–353 e10, 2017. ISSN 1878-1551 (Electronic) 1534-5807 (Linking). doi: 10.1016/j.devcel.2017.02.001.

17. H. Miyazawa, Y. Yamaguchi, Y. Sugiura, K. Honda, K. Kondo, F. Matsuda, T. Yamamoto, M. Suematsu, and M. Miura. Rewiring of embryonic glucose metabolism via suppression of pfk-1 and aldolase during mouse chorioallantoic branching. Development, 144(1):63–73, 2017. ISSN 1477-9129 (Electronic) 0950-1991 (Linking). doi: 10.1242/dev.138545.

18. D. Bhattacharya, A. P. Azambuja, and M. Simoes-Costa. Metabolic reprogramming promotes neural crest migration via yap/tead signaling. Dev Cell, 53(2):199–211 e6, 2020. ISSN 1878-1551 (Electronic) 1534-5807 (Linking). doi: 10.1016/j.devcel.2020.03.005.

19. N. J. Djabrayan, C. M. Smits, M. Krajnc, T. Stern, S. Yamada, W. C. Lemon, P. J. Keller, C. A. Rushlow, and S. Y. Shvartsman. Metabolic regulation of developmental cell cycles and zygotic transcription. Curr Biol, 29(7):1193–1198 e5, 2019. ISSN 1879-0445 (Electronic) 0960-9822 (Linking). doi: 10.1016/j.cub.2019.02.028.

20. J. Rodenfels, K. M. Neugebauer, and J. Howard. Heat oscillations driven by the embryonic cell cycle reveal the energetic costs of signaling. Dev Cell, 48(5):646–658 e6, 2019. ISSN 1878-1551 (Electronic) 1534-5807 (Linking). doi: 10.1016/j.devcel.2018.12.024.

21. F. Chi, M. S. Sharpley, R. Nagaraj, S. S. Roy, and U. Banerjee. Glycolysis-independent glucose metabolism distinguishes te from icm fate during mammalian embryogenesis. Dev Cell, 53(1):9–26 e4, 2020. ISSN 1878-1551 (Electronic) 1534-5807 (Linking). doi: 10.1016/j.devcel.2020.02.015.

22. A. Hubaud and O. Pourquie. Signalling dynamics in vertebrate segmentation. Nat Rev Mol Cell Biol, 15(11):709–21, 2014. ISSN 1471-0080 (Electronic) 1471-0072 (Linking). doi: 10.1038/nrm3891.

23. A. Aulehla, W. Wiegraebe, V. Baubet, M. B. Wahl, C. Deng, M. Taketo, M. Lewandoski, and O. Pourquie. A beta-catenin gradient links the clock and wavefront systems in mouse embryo segmentation. Nat Cell Biol, 10(2):186–93, 2008. ISSN 1476-4679 (Electronic) 1465-7392 (Linking). doi: 10.1038/ncb1679.

24. K. Yoshioka-Kobayashi, M. Matsumiya, Y. Niino, A. Isomura, H. Kori, A. Miyawaki, and R. Kageyama. Coupling delay controls synchronized oscillation in the segmentation clock. Nature, 580(7801):119–123, 2020. ISSN 1476-4687 (Electronic) 0028-0836 (Linking). doi: 10.1038/s41586-019-1882-z.

25. D. Soroldoni, D. J. Jorg, L. G. Morelli, D. L. Richmond, J. Schindelin, F. Julicher, and A. C. Oates. Genetic oscillations. a doppler effect in embryonic pattern formation. Science, 345(6193):222–5, 2014. ISSN 1095-9203 (Electronic) 0036-8075 (Linking). doi: 10.1126/science.1253089.

26. M. Matsuda, H. Hayashi, J. Garcia-Ojalvo, K. Yoshioka-Kobayashi, R. Kageyama, Y. Yamanaka, M. Ikeya, J. Toguchida, C. Alev, and M. Ebisuya. Species-specific segmentation clock periods are due to differential biochemical reaction speeds. Science, 369(6510): 1450–1455, 2020. ISSN 1095-9203 (Electronic) 0036-8075 (Linking). doi: 10.1126/science.aba7668.

27. M. Diaz-Cuadros, D. E. Wagner, C. Budjan, A. Hubaud, O. A. Tarazona, S. Donelly, Michaut, Z. Al Tanoury, K. Yoshioka-Kobayashi, Y. Niino, R. Kageyama, A. Miyawaki, J. Touboul, and O. Pourquie. In vitro characterization of the human segmentation clock. Nature, 580(7801):113–118, 2020. ISSN 1476-4687 (Electronic) 0028-0836 (Linking). doi: 10.1038/s41586-019-1885-9.

28. L. F. Chu, D. Mamott, Z. Ni, R. Bacher, C. Liu, S. Swanson, C. Kendziorski, R. Stewart, and J. A. Thomson. An in vitro human segmentation clock model derived from embryonic stem cells. Cell Rep, 28(9):2247–2255 e5, 2019. ISSN 2211-1247 (Electronic). doi: 10.1016/j.celrep.2019.07.090.

29. K. F. Sonnen, V. M. Lauschke, J. Uraji, H. J. Falk, Y. Petersen, M. C. Funk, M. Beaupeux, P. Francois, C. A. Merten, and A. Aulehla. Modulation of phase shift between wnt and notch signaling oscillations controls mesoderm segmentation. Cell, 172(5):1079–1090 e12, 2018. ISSN 1097-4172 (Electronic) 0092-8674 (Linking). doi: 10.1016/j.cell.2018.01.026.

30. M. Oginuma, Y. Harima, O. A. Tarazona, M. Diaz-Cuadros, A. Michaut, T. Ishitani, F. Xiong, and O. Pourquie. Intracellular ph controls wnt downstream of glycolysis in amniote embryos. Nature, 584(7819):98–101, 2020. ISSN 1476-4687 (Electronic) 0028-0836 (Linking). doi: 10.1038/s41586-020-2428-0.

31. Y. Niwa, Y. Masamizu, T. Liu, R. Nakayama, C. X. Deng, and R. Kageyama. The initiation and propagation of hes7 oscillation are cooperatively regulated by fgf and notch signaling in the somite segmentation clock. Dev Cell, 13(2):298–304, 2007. ISSN 1534-5807 (Print) 1534-5807 (Linking). doi: 10.1016/j.devcel.2007.07.013.

32. L. Wittler, E. H. Shin, P. Grote, A. Kispert, A. Beckers, A. Gossler, M. Werber, and B. G. Herrmann. Expression of msgn1 in the presomitic mesoderm is controlled by synergism of wnt signalling and tbx6. EMBO Rep, 8(8):784–9, 2007. ISSN 1469-221X (Print) 1469-221X (Linking). doi: 10.1038/sj.embor.7401030.

33. L. B. Tanner, A. G. Goglia, M. H. Wei, T. Sehgal, L. R. Parsons, J. O. Park, E. White, J. E. Toettcher, and J. D. Rabinowitz. Four key steps control glycolytic flux in mammalian cells. Cell Syst, 7(1):49–62 e8, 2018. ISSN 2405-4712 (Print) 2405-4712 (Linking). doi: 10.1016/j.cels.2018.06.003.

34. I. Mor, E. C. Cheung, and K. H. Vousden. Control of glycolysis through regulation of pfk1: old friends and recent additions. Cold Spring Harb Symp Quant Biol, 76:211–6, 2011. ISSN 1943-4456 (Electronic) 0091-7451 (Linking). doi: 10.1101/sqb.2011.76.010868.

35. A. Yalcin, B. F. Clem, A. Simmons, A. Lane, K. Nelson, A. L. Clem, E. Brock, D. Siow, B. Wattenberg, S. Telang, and J. Chesney. Nuclear targeting of 6-phosphofructo-2-kinase (pfkfb3) increases proliferation via cyclin-dependent kinases. J Biol Chem, 284(36):24223– 32, 2009. ISSN 0021-9258 (Print) 0021-9258 (Linking). doi: 10.1074/jbc.M109.016816.

36. A. O. Perantoni, O. Timofeeva, F. Naillat, C. Richman, S. Pajni-Underwood, C. Wilson, S. Vainio, L. F. Dove, and M. Lewandoski. Inactivation of fgf8 in early mesoderm reveals an essential role in kidney development. Development, 132(17):3859–71, 2005. ISSN 0950-1991 (Print) 0950-1991 (Linking). doi: 10.1242/dev.01945.

37. J. J. Wilde, J. R. Petersen, and L. Niswander. Genetic, epigenetic, and environmental contributions to neural tube closure. Annu Rev Genet, 48:583–611, 2014. ISSN 1545-2948 (Electronic) 0066-4197 (Linking). doi: 10.1146/annurev-genet-120213-092208.

38. J. A. Rivera-Perez, V. Jones, and P. P. Tam. Culture of whole mouse embryos at early postimplantation to organogenesis stages: developmental staging and methods. Methods Enzymol, 476:185–203, 2010. ISSN 1557-7988 (Electronic) 0076-6879 (Linking). doi: 10.1016/S0076-6879(10)76011-0.

39. M. B. Renfree, H. C. Hensleigh, and A. McLaren. Developmental changes in the composition and amount of mouse fetal fluids. J Embryol Exp Morphol, 33(2):435–46, 1975. ISSN 0022-0752 (Print) 0022-0752 (Linking).

40. H. J. Kwon, J. H. Rhim, I. S. Jang, G. E. Kim, S. C. Park, and E. J. Yeo. Activation of amp-activated protein kinase stimulates the nuclear localization of glyceraldehyde 3-phosphate dehydrogenase in human diploid fibroblasts. Exp Mol Med, 42(4):254–69, 2010. ISSN 2092-6413 (Electronic) 1226-3613 (Linking). doi: 10.3858/emm.2010.42.4.025.

41. H. Hu, A. Juvekar, C. A. Lyssiotis, E. C. Lien, J. G. Albeck, D. Oh, G. Varma, Y. P. Hung, S. Ullas, J. Lauring, P. Seth, M. R. Lundquist, D. R. Tolan, A. K. Grant, D. J. Needleman, J. M. Asara, L. C. Cantley, and G. M. Wulf. Phosphoinositide 3-kinase regulates glycolysis through mobilization of aldolase from the actin cytoskeleton. Cell, 164(3):433–46, 2016. ISSN 1097-4172 (Electronic) 0092-8674 (Linking). doi: 10.1016/j.cell.2015.12.042.

42. C. S. Zhang, S. A. Hawley, Y. Zong, M. Li, Z. Wang, A. Gray, T. Ma, J. Cui, J. W. Feng, M. Zhu, Y. Q. Wu, T. Y. Li, Z. Ye, S. Y. Lin, H. Yin, H. L. Piao, D. G. Hardie, and S. C. Lin. Fructose-1,6-bisphosphate and aldolase mediate glucose sensing by ampk. Nature, 548 (7665):112–116, 2017. ISSN 1476-4687 (Electronic) 0028-0836 (Linking). doi: 10.1038/nature23275.

43. K. Kochanowski, B. Volkmer, L. Gerosa, B. R. Haverkorn van Rijsewijk, A. Schmidt, and M. Heinemann. Functioning of a metabolic flux sensor in escherichia coli. Proc Natl Acad Sci U S A, 110(3):1130–5, 2013. ISSN 1091-6490 (Electronic) 0027-8424 (Linking). doi: 10.1073/pnas.1202582110.

44. L. Cai, B. M. Sutter, B. Li, and B. P. Tu. Acetyl-coa induces cell growth and proliferation by promoting the acetylation of histones at growth genes. Mol Cell, 42(4):426–37, 2011. ISSN 1097-4164 (Electronic) 1097-2765 (Linking). doi: 10.1016/j.molcel.2011.05.004.

45. C. Jang, L. Chen, and J. D. Rabinowitz. Metabolomics and isotope tracing. Cell, 173(4): 822–837, 2018. ISSN 1097-4172 (Electronic) 0092-8674 (Linking). doi: 10.1016/j.cell.2018.03.055.

46. O. Kotte, J. B. Zaugg, and M. Heinemann. Bacterial adaptation through distributed sensing of metabolic fluxes. Mol Syst Biol, 6:355, 2010. ISSN 1744-4292 (Electronic) 1744-4292 (Linking). doi: 10.1038/msb.2010.10.

47. N. Alva, R. Alva, and T. Carbonell. Fructose 1,6-bisphosphate: A summary of its cytoprotective mechanism. Curr Med Chem, 23(39):4396–4417, 2016. ISSN 1875-533X (Electronic) 0929-8673 (Linking). doi: 10.2174/0929867323666161014144250.

48. L. N. Duysens and J. Amesz. Fluorescence spectrophotometry of reduced phosphopyridine nucleotide in intact cells in the near-ultraviolet and visible region. Biochim Biophys Acta, 24 (1):19–26, 1957. ISSN 0006-3002 (Print) 0006-3002 (Linking). doi: 10.1016/0006-3002(57)90141-5.

49. B. Chance, R. W. Estabrook, and A. Ghosh. Damped sinusoidal oscillations of cytoplasmic reduced pyridine nucleotide in yeast cells. Proc Natl Acad Sci U S A, 51:1244–51, 1964. ISSN 0027-8424 (Print) 0027-8424 (Linking). doi: 10.1073/pnas.51.6.1244.

50. A. Goldbeter and M. J. Berridge. Biochemical Oscillations and Cellular Rhythms: The Molecular Bases of Periodic and Chaotic Behaviour. Cambridge University Press, 1996. doi: 10.1017/CBO9780511608193.

51. K. T. Pate, C. Stringari, S. Sprowl-Tanio, K. Wang, T. TeSlaa, N. P. Hoverter, M. M. McQuade, C. Garner, M. A. Digman, M. A. Teitell, R. A. Edwards, E. Gratton, and M. L. Waterman. Wnt signaling directs a metabolic program of glycolysis and angiogenesis in colon cancer. EMBO J, 33(13):1454–73, 2014. ISSN 1460-2075 (Electronic) 0261-4189 (Linking). doi: 10.15252/embj.201488598.

52. E. Enzo, G. Santinon, A. Pocaterra, M. Aragona, S. Bresolin, M. Forcato, D. Grifoni, A. Pession, F. Zanconato, G. Guzzo, S. Bicciato, and S. Dupont. Aerobic glycolysis tunes yap/taz transcriptional activity. EMBO J, 34(10):1349–70, 2015. ISSN 1460-2075 (Electronic) 0261-4189 (Linking). doi: 10.15252/embj.201490379.

53. M. Ciesla, J. Mierzejewska, M. Adamczyk, A. K. Farrants, and M. Boguta. Fructose bisphos-phate aldolase is involved in the control of rna polymerase iii-directed transcription. Biochim Biophys Acta, 1843(6):1103–10, 2014. ISSN 0006-3002 (Print) 0006-3002 (Linking). doi: 10.1016/j.bbamcr.2014.02.007.

54. Z. Ronai, R. Robinson, S. Rutberg, P. Lazarus, and M. Sardana. Aldolase-dna interactions in a sewa cell system. Biochim Biophys Acta, 1130(1):20–8, 1992. ISSN 0006-3002 (Print) 0006-3002 (Linking). doi: 10.1016/0167-4781(92)90456-a.

55. W. Yang, Y. Xia, H. Ji, Y. Zheng, J. Liang, W. Huang, X. Gao, K. Aldape, and Z. Lu. Nuclear pkm2 regulates beta-catenin transactivation upon egfr activation. Nature, 480(7375):118– 22, 2011. ISSN 1476-4687 (Electronic) 0028-0836 (Linking). doi: 10.1038/nature10598.

56. W. Yang, Y. Zheng, Y. Xia, H. Ji, X. Chen, F. Guo, C. A. Lyssiotis, K. Aldape, L. C. Cantley, and Z. Lu. Erk1/2-dependent phosphorylation and nuclear translocation of pkm2 promotes the warburg effect. Nat Cell Biol, 14(12):1295–304, 2012. ISSN 1476-4679 (Electronic) 1465-7392 (Linking). doi: 10.1038/ncb2629.

57. R. Nagaraj, M. S. Sharpley, F. Chi, D. Braas, Y. Zhou, R. Kim, A. T. Clark, and U. Banerjee. Nuclear localization of mitochondrial tca cycle enzymes as a critical step in mammalian zygotic genome activation. Cell, 168(1-2):210–223 e11, 2017. ISSN 1097-4172 (Electronic) 0092-8674 (Linking). doi: 10.1016/j.cell.2016.12.026.

58. E. Kafkia, A. Andres-Pons, K. Ganter, M. Seiler, P. Jouhten, F. Pereira, J. B. Zaugg, C. Lancrin, M. Beck, and K. R. Patil. Operation of a TCA cycle subnetwork in the mammalian nucleus. bioRxiv, 2020. doi: 10.1101/2020.11.22.393413.

59. K. De Bock, M. Georgiadou, S. Schoors, A. Kuchnio, B. W. Wong, A. R. Cantelmo, Quaegebeur, B. Ghesquiere, S. Cauwenberghs, G. Eelen, L. K. Phng, I. Betz, B. Tembuyser, K. Brepoels, J. Welti, I. Geudens, I. Segura, B. Cruys, F. Bifari, I. Decimo, R. Blanco, S. Wyns, J. Vangindertael, S. Rocha, R. T. Collins, S. Munck, D. Daelemans, H. Imamura, R. Devlieger, M. Rider, P. P. Van Veldhoven, F. Schuit, R. Bartrons, J. Hofkens, P. Fraisl, S. Telang, R. J. Deberardinis, L. Schoonjans, S. Vinckier, J. Chesney, H. Gerhardt, M. Dewerchin, and P. Carmeliet. Role of pfkfb3-driven glycolysis in vessel sprouting. Cell, 154(3):651–63, 2013. ISSN 1097-4172 (Electronic) 0092-8674 (Linking). doi: 10.1016/j.cell.2013.06.037.

60. S. Jang, J. C. Nelson, E. G. Bend, L. Rodriguez-Laureano, F. G. Tueros, L. Cartagenova, K. Underwood, E. M. Jorgensen, and D. A. Colon-Ramos. Glycolytic enzymes localize to synapses under energy stress to support synaptic function. Neuron, 90(2):278–91, 2016. ISSN 1097-4199 (Electronic) 0896-6273 (Linking). doi: 10.1016/j.neuron.2016.03.011.

61. K. W. Ryu, T. Nandu, J. Kim, S. Challa, R. J. DeBerardinis, and W. L. Kraus. Metabolic regulation of transcription through compartmentalized nad(+) biosynthesis. Science, 360 (6389), 2018. ISSN 1095-9203 (Electronic) 0036-8075 (Linking). doi: 10.1126/science.aan5780.

62. I. Piazza, K. Kochanowski, V. Cappelletti, T. Fuhrer, E. Noor, U. Sauer, and P. Picotti. A map of protein-metabolite interactions reveals principles of chemical communication. Cell, 172 (1-2):358–372 e23, 2018. ISSN 1097-4172 (Electronic) 0092-8674 (Linking). doi: 10.1016/j.cell.2017.12.006.

63. J. E. Lindsley and J. Rutter. Whence cometh the allosterome? Proc Natl Acad Sci U S A, 103(28):10533–5, 2006. ISSN 0027-8424 (Print) 0027-8424 (Linking). doi: 10.1073/pnas.0604452103.

64. M. M. Savitski, F. B. Reinhard, H. Franken, T. Werner, M. F. Savitski, D. Eberhard, D. Martinez Molina, R. Jafari, R. B. Dovega, S. Klaeger, B. Kuster, P. Nordlund, M. Bantscheff, and G. Drewes. Tracking cancer drugs in living cells by thermal profiling of the proteome. Science, 346(6205):1255784, 2014. ISSN 1095-9203 (Electronic) 0036-8075 (Linking). doi: 10.1126/science.1255784.

65. L. C. Schulz. The dutch hunger winter and the developmental origins of health and disease. Proc Natl Acad Sci U S A, 107(39):16757–8, 2010. ISSN 1091-6490 (Electronic) 0027-8424 (Linking). doi: 10.1073/pnas.1012911107.

66. D. B. Sparrow, G. Chapman, A. J. Smith, M. Z. Mattar, J. A. Major, V. C. O’Reilly, Y. Saga, E. H. Zackai, J. P. Dormans, B. A. Alman, L. McGregor, R. Kageyama, K. Kusumi, and S. L. Dunwoodie. A mechanism for gene-environment interaction in the etiology of congenital scoliosis. Cell, 149(2):295–306, 2012. ISSN 1097-4172 (Electronic) 0092-8674 (Linking). doi: 10.1016/j.cell.2012.02.054.

67. T. W. Holstein. The evolution of the wnt pathway. Cold Spring Harb Perspect Biol, 4 (7):a007922, 2012. ISSN 1943-0264 (Electronic) 1943-0264 (Linking). doi: 10.1101/cshperspect.a007922.

68. L. Madisen, T. A. Zwingman, S. M. Sunkin, S. W. Oh, H. A. Zariwala, H. Gu, L. L. Ng, R. D. Palmiter, M. J. Hawrylycz, A. R. Jones, E. S. Lein, and H. Zeng. A robust and high-throughput cre reporting and characterization system for the whole mouse brain. Nat Neurosci, 13(1):133–40, 2010. ISSN 1546-1726 (Electronic) 1097-6256 (Linking). doi: 10.1038/nn.2467.

69. S. H. Tang, F. J. Silva, W. M. Tsark, and J. R. Mann. A cre/loxp-deleter transgenic line in mouse strain 129s1/svimj. Genesis, 32(3):199–202, 2002. ISSN 1526-954X (Print) 1526-954X (Linking). doi: 10.1002/gene.10030.

70. V. M. Lauschke, C. D. Tsiairis, P. Francois, and A. Aulehla. Scaling of embryonic patterning based on phase-gradient encoding. Nature, 493(7430):101–5, 2013. ISSN 1476-4687 (Electronic) 0028-0836 (Linking). doi: 10.1038/nature11804.

71. J. Schindelin, I. Arganda-Carreras, E. Frise, V. Kaynig, M. Longair, T. Pietzsch, S. Preibisch, C. Rueden, S. Saalfeld, B. Schmid, J. Y. Tinevez, D. J. White, V. Hartenstein, K. Eliceiri, P. Tomancak, and A. Cardona. Fiji: an open-source platform for biological-image analysis. Nat Methods, 9(7):676–82, 2012. ISSN 1548-7105 (Electronic) 1548-7091 (Linking). doi: 10.1038/nmeth.2019.

72. H. H. Kanani and M. I. Klapa. Data correction strategy for metabolomics analysis using gas chromatography-mass spectrometry. Metab Eng, 9(1):39–51, 2007. ISSN 1096-7176 (Print) 1096-7176 (Linking). doi: 10.1016/j.ymben.2006.08.001.

73. S. Blasche, Y. Kim, R. A. T. Mars, D. Machado, M. Maansson, E. Kafkia, A. Milanese, G. Zeller, B. Teusink, J. Nielsen, V. Benes, R. Neves, U. Sauer, and K. R. Patil. Metabolic cooperation and spatiotemporal niche partitioning in a kefir microbial community. Nat Microbiol, 6(2):196–208, 2021. ISSN 2058-5276 (Electronic) 2058-5276 (Linking). doi: 10.1038/s41564-020-00816-5.

74. E. Melamud, L. Vastag, and J. D. Rabinowitz. Metabolomic analysis and visualization engine for lc-ms data. Anal Chem, 82(23):9818–26, 2010. ISSN 1520-6882 (Electronic) 0003-2700 (Linking). doi: 10.1021/ac1021166.

75. C. S. Hughes, S. Foehr, D. A. Garfield, E. E. Furlong, L. M. Steinmetz, and J. Krijgsveld. Ultrasensitive proteome analysis using paramagnetic bead technology. Mol Syst Biol, 10: 757, 2014. ISSN 1744-4292 (Electronic) 1744-4292 (Linking). doi: 10.15252/msb.20145625.

76. Y. Perez-Riverol, A. Csordas, J. Bai, M. Bernal-Llinares, S. Hewapathirana, D. J. Kundu, Inuganti, J. Griss, G. Mayer, M. Eisenacher, E. Perez, J. Uszkoreit, J. Pfeuffer, T. Sachsenberg, S. Yilmaz, S. Tiwary, J. Cox, E. Audain, M. Walzer, A. F. Jarnuczak, T. Ternent, A. Brazma, and J. A. Vizcaino. The pride database and related tools and resources in 2019: improving support for quantification data. Nucleic Acids Res, 47(D1):D442–D450, 2019. ISSN 1362-4962 (Electronic) 0305-1048 (Linking). doi: 10.1093/nar/gky1106.

77. H. Franken, T. Mathieson, D. Childs, G. M. Sweetman, T. Werner, I. Togel, C. Doce, S. Gade, M. Bantscheff, G. Drewes, F. B. Reinhard, W. Huber, and M. M. Savitski. Thermal proteome profiling for unbiased identification of direct and indirect drug targets using multiplexed quantitative mass spectrometry. Nat Protoc, 10(10):1567–93, 2015. ISSN 1750-2799 (Electronic) 1750-2799 (Linking). doi: 10.1038/nprot.2015.101.

78. W. Huber, A. von Heydebreck, H. Sultmann, A. Poustka, and M. Vingron. Variance stabilization applied to microarray data calibration and to the quantification of differential expression. Bioinformatics, 18 Suppl 1:S96–104, 2002. ISSN 1367-4803 (Print) 1367-4803 (Linking). doi: 10.1093/bioinformatics/18.suppl_1.s96.

79. M. E. Ritchie, B. Phipson, D. Wu, Y. Hu, C. W. Law, W. Shi, and G. K. Smyth. limma powers differential expression analyses for rna-sequencing and microarray studies. Nucleic Acids Res, 43(7):e47, 2015. ISSN 1362-4962 (Electronic) 0305-1048 (Linking). doi: 10.1093/nar/gkv007.

80. V. G. Tusher, R. Tibshirani, and G. Chu. Significance analysis of microarrays applied to the ionizing radiation response. Proc Natl Acad Sci U S A, 98(9):5116–21, 2001. ISSN 0027-8424 (Print) 0027-8424 (Linking). doi: 10.1073/pnas.091062498.

81. C. R. Harris, K. J. Millman, S. J. van der Walt, R. Gommers, P. Virtanen, D. Cournapeau, E. Wieser, J. Taylor, S. Berg, N. J. Smith, R. Kern, M. Picus, S. Hoyer, M. H. van Kerkwijk, M. Brett, A. Haldane, J. F. Del Rio, M. Wiebe, P. Peterson, P. Gerard-Marchant, K. Sheppard, T. Reddy, W. Weckesser, H. Abbasi, C. Gohlke, and T. E. Oliphant. Array programming with numpy. Nature, 585(7825):357–362, 2020. ISSN 1476-4687 (Electronic) 0028-0836 (Linking). doi: 10.1038/s41586-020-2649-2.

82. Wes McKinney. Data Structures for Statistical Computing in Python. In Stéfan van der Walt and Jarrod Millman, editors, Proceedings of the 9th Python in Science Conference, pages 56–61, 2010. doi: 10.25080/Majora-92bf1922-00a.

83. P. Virtanen, R. Gommers, T. E. Oliphant, M. Haberland, T. Reddy, D. Cournapeau, E. Burovski, P. Peterson, W. Weckesser, J. Bright, S. J. van der Walt, M. Brett, J. Wilson, K. J. Millman, N. Mayorov, A. R. J. Nelson, E. Jones, R. Kern, E. Larson, C. J. Carey, I. Polat, Y. Feng, E. W. Moore, J. VanderPlas, D. Laxalde, J. Perktold, R. Cimrman, I. Henriksen, E. A. Quintero, C. R. Harris, A. M. Archibald, A. H. Ribeiro, F. Pedregosa, P. van Mulbregt, and Contributors SciPy. Scipy 1.0: fundamental algorithms for scientific computing in python. Nat Methods, 17(3):261–272, 2020. ISSN 1548-7105 (Electronic) 1548-7091 (Linking). doi: 10.1038/s41592-019-0686-2.

84. J. D. Hunter. Matplotlib: A 2d graphics environment. Computing in Science & Engineering, 9(3):90–95, 2007. doi: 10.1109/MCSE.2007.55.

